# Glutathione supports lipid abundance *in vivo*

**DOI:** 10.1101/2023.02.10.524960

**Authors:** Gloria Asantewaa, Emily T. Tuttle, Nathan P. Ward, Yun Pyo Kang, Yumi Kim, Madeline E. Kavanagh, Nomeda Girnius, Ying Chen, Renae Duncan, Katherine Rodriguez, Fabio Hecht, Marco Zocchi, Leonid Smorodintsev-Schiller, TashJaé Q. Scales, Kira Taylor, Fatemeh Alimohammadi, Zachary R Sechrist, Diana Agostini-Vulaj, Xenia L. Schafer, Hayley Chang, Zachary Smith, Thomas N. O’Connor, Sarah Whelan, Laura M. Selfors, Jett Crowdis, G. Kenneth Gray, Roderick T. Bronson, Dirk Brenner, Alessandro Rufini, Robert T. Dirksen, Aram F. Hezel, Aaron R. Huber, Josh Munger, Benjamin F. Cravatt, Vasilis Vasiliou, Calvin L Cole, Gina M. DeNicola, Isaac S. Harris

**Affiliations:** Department of Biochemistry and Biophysics, University of Rochester Medical Center, Rochester, NY, USA, 14642; Department of Biomedical Genetics, University of Rochester Medical Center, Rochester, NY, USA, 14642; Wilmot Cancer Institute, University of Rochester Medical Center, Rochester, NY, USA, 14642; Department of Metabolism and Physiology, Moffitt Cancer Center and Research Institute, Tampa, FL, USA, 33612; Department of Chemistry and The Skaggs Institute for Chemical Biology, The Scripps Research Institute, La Jolla, CA, USA, 92037; Department of Cell Biology, Harvard Medical School, Boston, MA, USA, 02115; Department of Environmental Health Sciences, Yale School of Public Health, New Haven, CT, USA, 06520; Department of Pharmacology and Physiology, University of Rochester, Rochester, NY, USA, 14642; Department of Surgery and Center for Musculoskeletal Research, University of Rochester Medical Center, Rochester, NY, USA, 14642; Department of Pathology and Laboratory Medicine, University of Rochester Medical Center, Rochester, NY, USA, 14642; Leicester Cancer Research Centre, University of Leicester, Leicester, LE2 7LX, UK; Experimental & Molecular Immunology, Department of Infection and Immunity, Luxembourg Institute of Health, 29 Rue Henri Koch, Esch-sur-Alzette, Luxembourg; Odense Research Center for Anaphylaxis (ORCA), Department of Dermatology and Allergy Center, Odense University Hospital, University of Southern Denmark, Odense, Denmark; Dipartimento di Bioscienze, Università degli Studi di Milano, Via Celoria 26, 20133, Milan, Italy

**Keywords:** glutathione, oxidative stress, NRF2, liver, lipids.

## Abstract

Cells rely on antioxidants to survive. The most abundant antioxidant is glutathione (GSH). The synthesis of GSH is non-redundantly controlled by the glutamate-cysteine ligase catalytic subunit (GCLC). GSH imbalance is implicated in many diseases, but the requirement for GSH in adult tissues is unclear. To interrogate this, we developed a series of *in vivo* models to induce *Gclc* deletion in adult animals. We find that GSH is essential to lipid abundance *in vivo*. GSH levels are reported to be highest in liver tissue, which is also a hub for lipid production. While the loss of GSH did not cause liver failure, it decreased lipogenic enzyme expression, circulating triglyceride levels, and fat stores. Mechanistically, we found that GSH promotes lipid abundance by repressing NRF2, a transcription factor induced by oxidative stress. These studies identify GSH as a fulcrum in the liver’s balance of redox buffering and triglyceride production.

## INTRODUCTION

Fundamental cellular processes, including the electron transport chain (ETC) in the mitochondria and protein folding in the ER, produce reactive oxygen species (ROS)^1^. The function of ROS can be either beneficial or damaging, depending on its cellular concentration and localization^2–4^. Antioxidative systems, including metabolites and enzymes, scavenge ROS. Disparities between antioxidant activity and ROS levels can lead to oxidative stress. Studies have shown impaired antioxidant functions connected to the onset and progression of numerous diseases^5^. Many of these studies, however, are correlative. Notably, the roles of antioxidants in normal physiology are poorly understood, largely preventing our ability to modulate antioxidants to either prevent or treat disease.

Glutathione (GSH) is the most abundant antioxidant in the body^6^ and one of the most concentrated metabolites in the cell^7^. Unlike most abundantly synthesized molecules, such as ATP or DNA, the production of GSH is controlled at the enzymatic level by a holoenzyme, glutamate-cysteine ligase (GCL), a heterodimer composed of a catalytic subunit (GCLC), and a modifier subunit (GCLM). Tissue-specific Gclc KO in mice has identified important functions for GSH in physiological settings^8–13^. However, the impact of GSH across all the tissues in adult animals is unknown because mice that lack GCLC undergo an early embryonic lethality^14–16^. The noncatalytic modifier subunit GCLM aids GCLC in GSH synthesis. Gclm KO mice are viable^17, 18^, having 15-40% of normal tissue GSH levels due to GCLC expression. Using chemical inhibitors to block GCLC in animals leads to inconsistent and incomplete loss of GSH across tissues^19^. Finally, many studies have interrogated the role of requisite amino acids (i.e., cysteine, glutamate, and glycine) or metabolic cofactors that regenerate GSH (i.e., NADPH)^20–25^ in GSH synthesis. Still, these molecules also have multiple GSH-independent functions, which can potentially confound interpretations. Thus, while GSH was discovered more than a century ago^26^, our understanding of its physiological contributions remains incomplete.

The liver is a significant source of triglyceride production in the body^27^. It is also a primary site of drug detoxification, which relies heavily on GSH availability^28^. Indeed, GSH is reported to be produced in the liver at levels higher than any other tissue in the body^29^. Whether this GSH store influences other liver functions (i.e., de novo lipogenesis) is unclear. Mice born with liver-specific deletion of *Gclc* (using a *Gclc*^f/f^ Albumin-Cre mouse strain) undergo liver failure and die shortly after birth^30^. Thus, the role of GSH synthesis in the liver and how this impacts the physiology of adult animals has yet to be fully elucidated.

The development of inducible conditional knockout mouse models has revealed several aspects of tissue homeostasis in animals^31–33^. We used this strategy to study the function of GSH *in vivo*. Following whole-body deletion of *Gclc*, adult mice rapidly lost weight, fat depots, and circulating triglycerides. We found that liver-specific GCLC expression in adult mice was not responsible for maintaining liver survival but instead sustained the expression of lipogenic enzymes and the abundance of triglycerides and fat depots in the body. Finally, we showed that GSH synthesis in the liver suppresses the activity of the antioxidant transcription factor nuclear factor erythroid 2-related factor 2 (NFE2L2, also known as NRF2). By restraining NRF2 activity, GSH synthesis facilitates the expression of lipogenic enzymes and enables the supply of triglycerides into circulation. Together, our work sheds light on the core functions of GSH synthesis *in vivo,* where we find it necessary for sustaining lipid abundance.

## RESULTS

### GSH synthesis is required for the survival of adult animals

To better understand the importance of GSH in adult mammals, we developed an inducible, whole-body Gclc KO mouse by breeding the *Gclc*^f/f^ mouse strain^30^ with the Rosa26-CreERT2 mouse strain^34^. *Gclc*^f/f^ Rosa26-CreERT2 mice (hereafter referred to as Gclc KO mice) were aged to adulthood (>12 weeks old) before being given tamoxifen (i.p. injection of 160 mg/kg tamoxifen for five consecutive days, as previously described^33^) to induce Cre recombinase activity and *Gclc* excision **(Figure 1A)**. *Gclc*^f/f^ mice (hereafter described as Gclc WT) were also treated with tamoxifen to control for potential confounding effects associated with tamoxifen treatment. Approximately 12-15 days following tamoxifen treatment, mouse tissues were analyzed. A robust decrease in *Gclc* mRNA and protein **(Figure 1B and 1C)** and GSH levels **(Figure 1D)** were observed across most tissues in Gclc KO mice compared with Gclc WT mice. Notably, the induced depletion of GSH in organs of adult animals was greater than previously observed depletions using pharmacologic approaches^19^. Notably, the brain had the lowest reduction in *Gclc* mRNA abundance **(Figure 1B)**. Although tamoxifen and its metabolites (i.e., 4-hydroxytamoxifen) can cross the blood-brain barrier, these compounds have reduced bioavailability in the brain compared to other tissues^35^; thus, potentially accounting for the lack of effective deletion observed. Deletion of Gclc caused a progressive and dramatic weight loss in mice, resulting in humane endpoints **(Figure 1E and 1F)**. A maximal deletion of *Gclc* was required to cause weight loss in mice, as heterozygous deletion did not cause mice to lose weight **(Figure S1A-S1C)**, suggesting that even minimal production of GSH can maintain homeostatic processes in the body. These phenotypes were not attributed to tamoxifen treatment, as treatment of control mice (*Gclc*^f/w^ mice) with tamoxifen failed to cause any weight changes **(Figure S1D)**. Further, no difference between biological sexes was observed in Gclc KO mice **(Figure S1E)**. Interestingly, the weight loss in Gclc KO mice was not associated with a corresponding loss in appetite, as no change in food consumption was observed **(Figure 1G)**. This suggests that the loss of GSH was specifically causing weight loss rather than a lack of food intake.

**Figure 1.**
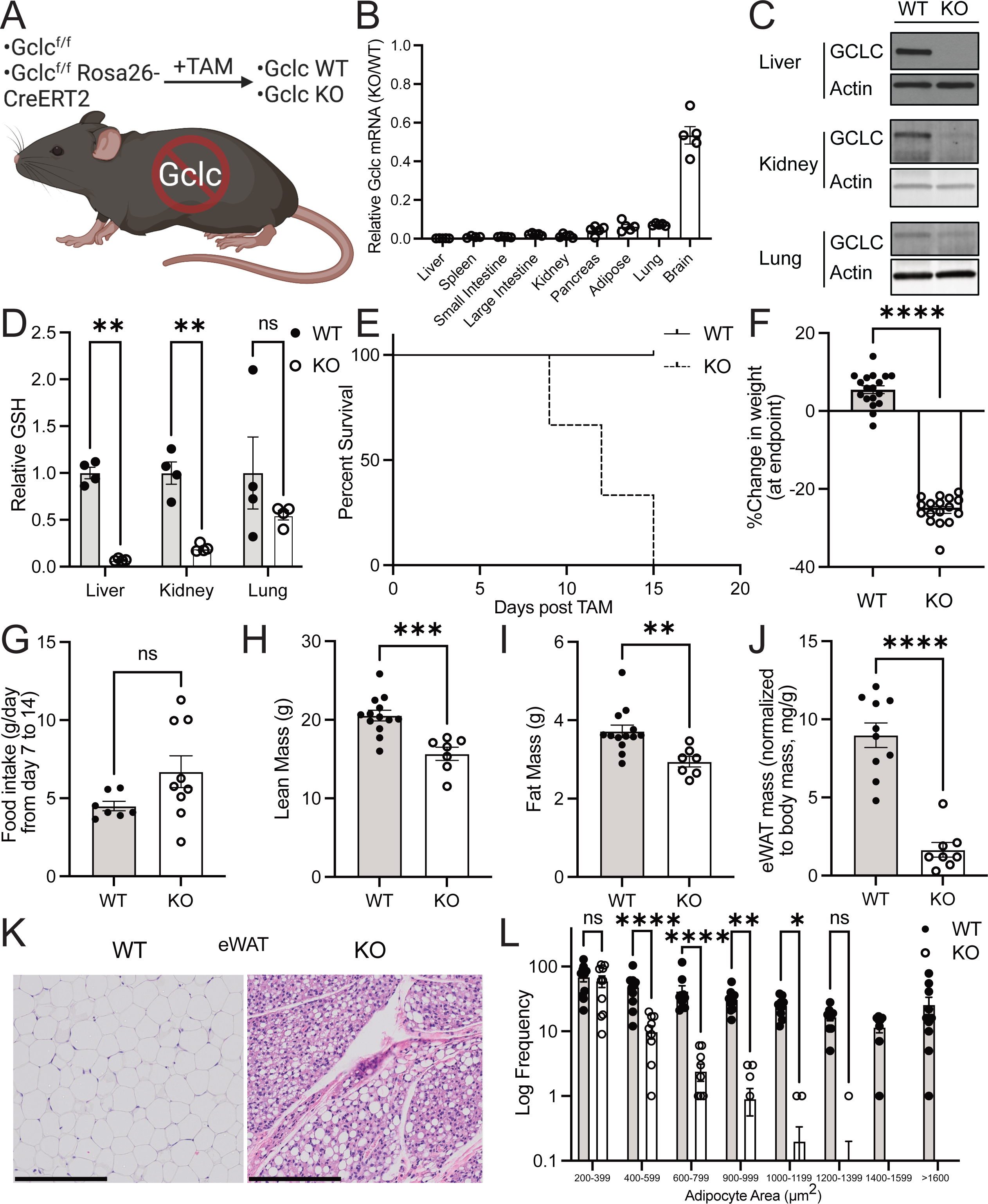
GSH synthesis is required for the survival of adult animals. (A) Schematic of the whole-body inducible Gclc knockout mouse model. (B) Relative expression of *Gclc* mRNA in tissues from the KO (n=4-5) compared to WT (n=5) mice 12-15 days following tamoxifen administration. Expression levels were normalized to the expression of the reference gene Rps9. (C) Representative immunoblot analysis of GCLC protein in the liver, kidney, and lung of WT and KO mice 12-15 days following tamoxifen administration. (D) Relative GSH abundance in the liver, kidney, and lungs of WT (n = 4) and KO (n = 4) mice 12-15 days following tamoxifen administration. (E) Percent survival of the WT (n = 9) and KO (n = 11) mice following tamoxifen treatment. Loss of greater than 20% body weight resulted in a humane endpoint for mice. (F) Percent change in body weight in WT (n=18) and KO (n=17) mice at death. Initial weight measurements were collected on Day 0 of tamoxifen administration, and final weight measurements were collected at endpoints. (G) Consumption of food by WT (n=7) and KO (n=5) mice over time. (H-I) Lean mass (H) and fat mass (I) in WT (n=13) and KO (n=7) mice as determined by dual-energy X-ray absorptiometry (DEXA) analysis on Day 14 post-tamoxifen injection. (J) Epididymal white adipose tissue (eWAT) mass normalized to body mass for WT (n=13) and KO (n=7) mice at endpoints. (K) Representative images from hematoxylin-eosin (H&E) staining of the epididymal fat from WT and KO mice. Tissues were collected 12-15 days following the tamoxifen injection (scale bars = 200 µm). (L) Adipocyte area measurements for WT and KO mice as quantified from representative H&E images with the Adiposoft plugin in ImageJ. N = 10 images for each genotype. Data are shown as mean ±SEM. A one-way ANOVA with subsequent Tukey’s multiple comparisons test was used in (D) and (L), and an unpaired two-tailed t-test was used in (F-J) to determine statistical significance. ns = not significant, * P value< 0.05, ** P value< 0.01, *** P value< 0.001, **** P value< 0.0001.

To better understand the cause of weight loss in GSH-depleted animals, we examined muscle and fat tissue from Gclc KO mice. Both lean and fat mass were decreased in Gclc KO mice compared to wild-type mice **(Figure 1H-1I)**. While muscle tissue from Gclc KO mice was lower in mass, it did not display histological hallmarks indicative of muscle damage, such as the increased prevalence of centrally-nucleated myofibers **(Figure S1F)**. **Quantifying and normalizing** organ masses to body mass revealed that no organs or tissues were lower in relative size except for fat depots **(Figure 1J and S1G-S1H)**. This included epididymal and inguinal white adipose tissue (eWAT and iWAT), brown adipose tissue (BAT), and mammary gland tissue. Histological analysis revealed that adipocytes in eWAT from Gclc KO mice were much smaller than Gclc WT mice, with a drastic reduction in the fat reservoir compartments and an increase in the extracellular matter **(Figure 1K-1L)**. These findings demonstrate that the production of GSH is required to maintain adipose tissue in adult animals.

### Loss of GSH synthesis induces transcription of NRF2 target genes and repression of lipogenic genes

GSH synthesis is required for numerous enzymatic processes across a range of cell types in the body. It is also a co-factor for GPX4-mediated lipid detoxification^36^, and GPX4 activity is necessary for the survival of several tissues, including the liver^37^ and kidney^38^. Further, the liver is reported to be one of the largest producers of GSH^29^. In line with previous findings, we show that the liver contained the highest levels of GSH compared to other organs **(Figure 2A)**. Surprisingly, Gclc KO mice showed no signs of liver failure, both by histology and serum biochemistry markers **(Figure 2B and S2A)**. While liver damage markers were increased in the serum of Gclc KO mice, their levels fell within the physiological range reported in healthy mice^39^. Indeed, other tissues from the Gclc KO mice, including the kidney, pancreas, and spleen, were also histologically similar to tissues from the Gclc WT mice **(Figure S2B-S2D)**. This suggests that, compared to the requirement for GSH synthesis in embryonic development^14–16^, GSH synthesis is dispensable to several organs in adult animals.

**Figure 2.**
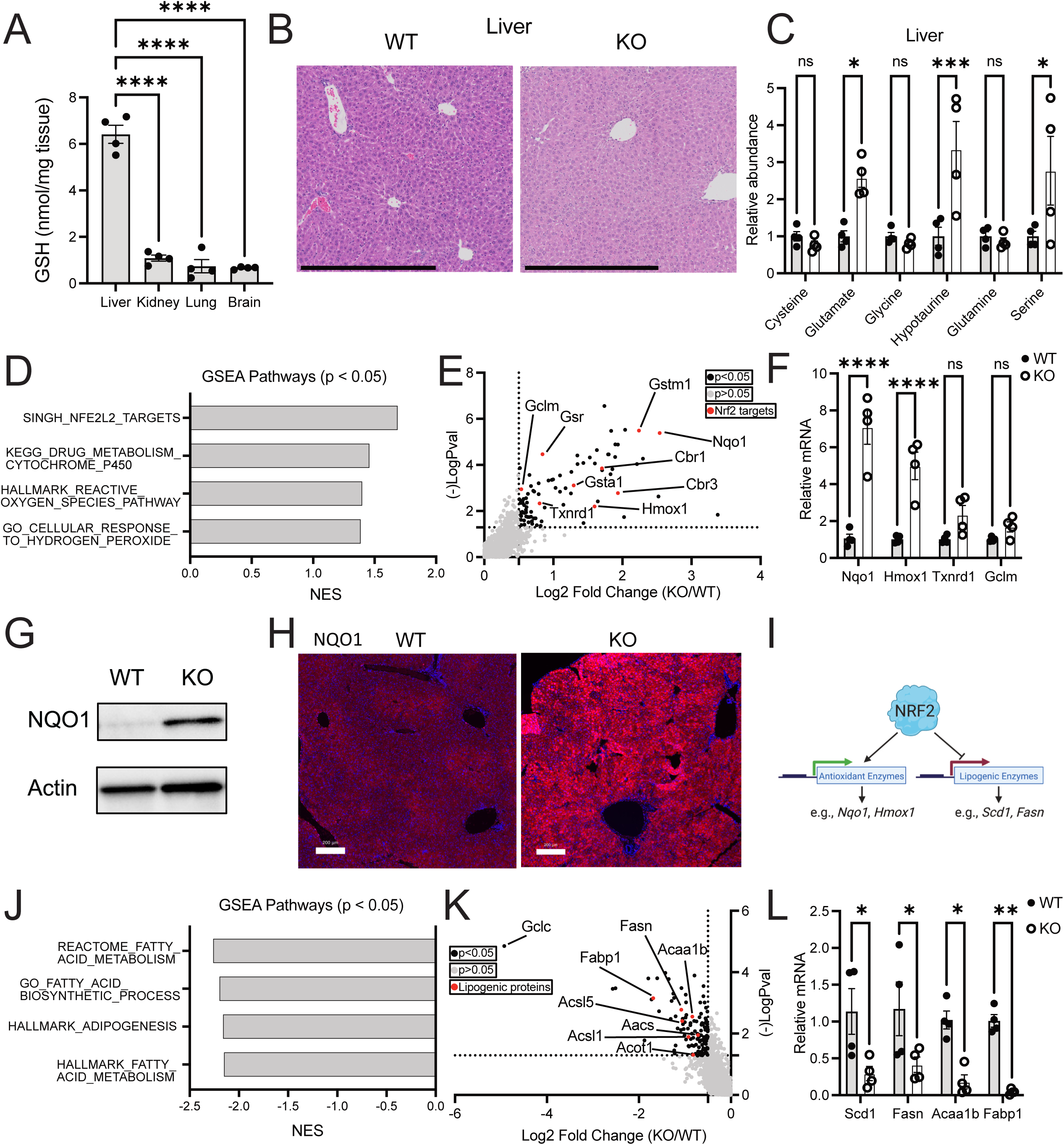
Liver tissue from Gclc KO mice has induced NRF2 target genes and repressed lipogenic gene expression. (A) GSH quantity (normalized to mg tissue weight) in the liver, kidney, lung, and brain from Gclc WT mice (n=4). (B) Representative H&E-stained histochemical images of the liver in WT and KO mice. Scale bars = 500 µm. (C) Relative abundance of GSH precursors and their related metabolites in the serum of WT (n=4) and KO (n=4) mice. (D) Gene Set Enrichment Analysis (GSEA) of oxidative stress-related pathways in the liver of KO (n=4) compared to WT (n=4) mice. (E) Proteomic analysis of upregulated liver proteins in KO (n=3) compared to WT (n=3) mice. Black data points = proteins with a p <0.05 and log2 fold change > 2. Red data points = annotated NRF2 target proteins. (F) Relative mRNA expression of annotated NRF2 target genes in the liver of WT (n=4) and KO (n=4) mice. Expression levels were normalized to the expression of the reference gene Rps9. (G) Representative immunoblot analysis of Nqo1 in the liver of WT and KO mice. (H) Representative immunofluorescence images of the liver from WT and KO mice stained with an antibody against NQO1. Scale bars = 200 µm. (I) Schematic of the proposed mechanism of NRF2-dependent repression of lipogenic gene expression. Data are shown as mean ±SEM. A one-way ANOVA with subsequent Tukey’s multiple comparisons test was used in (D) and (L), and an unpaired two-tailed t-test was used in (F-J) to determine statistical significance. ns = not significant, * P value < 0.05, ** P value < 0.01, *** P value < 0.001, **** P value < 0.0001.

The amino acids required to synthesize the GSH (cysteine, glutamate, and glycine) are also essential to many cellular processes, ranging from energy generation to protein synthesis^40^. Further, these precursors can be converted into other metabolites. Cysteine can be metabolized into cysteine sulphinate (via CDO1), which then can produce hypotaurine (via CSAD)^41^. Glycine can be interchanged with serine^42^, and glutamate can be converted to glutamine^43^. To determine if GSH precursors accumulate and are possibly rerouted upon halting GSH synthesis *in vivo*, we performed targeted metabolomics on tissues from Gclc WT and KO mice. In liver tissue from Gclc KO mice, we observed an accumulation of glutamate but no difference in cysteine and glycine levels **(Figure 2C)**. We hypothesized that the lack of accumulated cysteine and glycine could be due to these metabolites entering alternative metabolic pathways. Indeed, we found hypotaurine and serine levels increased in the liver of Gclc KO mice **(Figure 2C)**. These data suggest that in the absence of GSH synthesis, the liver accumulates GSH precursors and can re-route them into alternative metabolic pathways.

We hypothesized that, in addition to metabolic alterations, other changes were occurring in the liver of Gclc KO mice, which could explain the lack of damage in the tissue. KEAP1 is oxidized and inactivated upon oxidative stress, stabilizing the antioxidant transcription factor NRF2^44^. Hepatocytes induce *Nrf2* and reprogram their antioxidant pathways to deal with oxidative stress^45, 46^. RNA-seq and subsequent gene set enrichment analysis (GSEA) revealed that the liver tissue from Gclc KO mice was enriched in several gene signatures associated with stress responses, including those related to the activation of NRF2 **(Figure 2D and Table S1)**. Consistent with the transcriptomic findings, proteomic analysis of liver from Gclc KO mice revealed that most significantly elevated proteins in Gclc KO liver were encoded by NRF2 target genes, including the canonical NRF2 target NQO1 (NAD(P)H Quinone Dehydrogenase 1) **(Figure 2E-2H and Table S2)**. This stress response was not limited to the liver but instead observed across multiple tissues from Gclc KO mice, including the kidney and colon **(Figure S2E)**. These results suggest that upon depletion of GSH, tissues undergo reprogramming to hyperactivate NRF2, possibly as a mechanism to upregulate adjacent antioxidant pathways and buffer oxidative stress induced upon GSH depletion.

Activation of NRF2 is reported to coincide with the repression of enzymes involved in adipogenesis **(Figure 2I)**^47, 48^. In addition to increased transcription of NRF2 target genes, livers from Gclc KO mice had lower expression of lipogenic genes and proteins **(Figure 2J-2L, S2F and Table S1)**, such as stearoyl-Coenzyme A desaturase 1 (Scd1), a key enzyme in adipogenesis. The co-occurrence of high NRF2 target genes with low lipogenic genes was shared by some tissues but not all **(Figure S2G)**. These results indicate that following GSH depletion, lipogenic gene expression is repressed. The extent to which this impacted lipogenic products (i.e., fatty acids and triglycerides) and whether this was connected to NRF2 activation was unclear.

### GSH synthesis is required to maintain circulating triglycerides

Systemic depletion of GSH disproportionally impacted adipose tissue mass. Multiple tissues showed lower lipogenic enzyme expression, suggesting lipid production might also be impaired. To examine this, we performed lipidomics on several tissues from Gclc KO mice **(Figure 3A-3B, S3A-S3B, and Table S3)**. We found that triglycerides were among the most significantly depleted lipid species across tissues from Gclc KO mice. Since this appeared to be a global trend, we hypothesized that circulating triglycerides were depleted in Gclc KO mice. Indeed, lipidomics revealed that nearly all the significantly depleted lipid species in serum from Gclc KO mice were triglycerides **(Figure 3C and Table S3)**. This phenotype was confirmed by targeted measurement of triglycerides in serum **(Figure 3D)**. Together, these data demonstrate that the abundance of triglycerides in Gclc KO is lower, especially in circulating pools.

**Figure 3:**
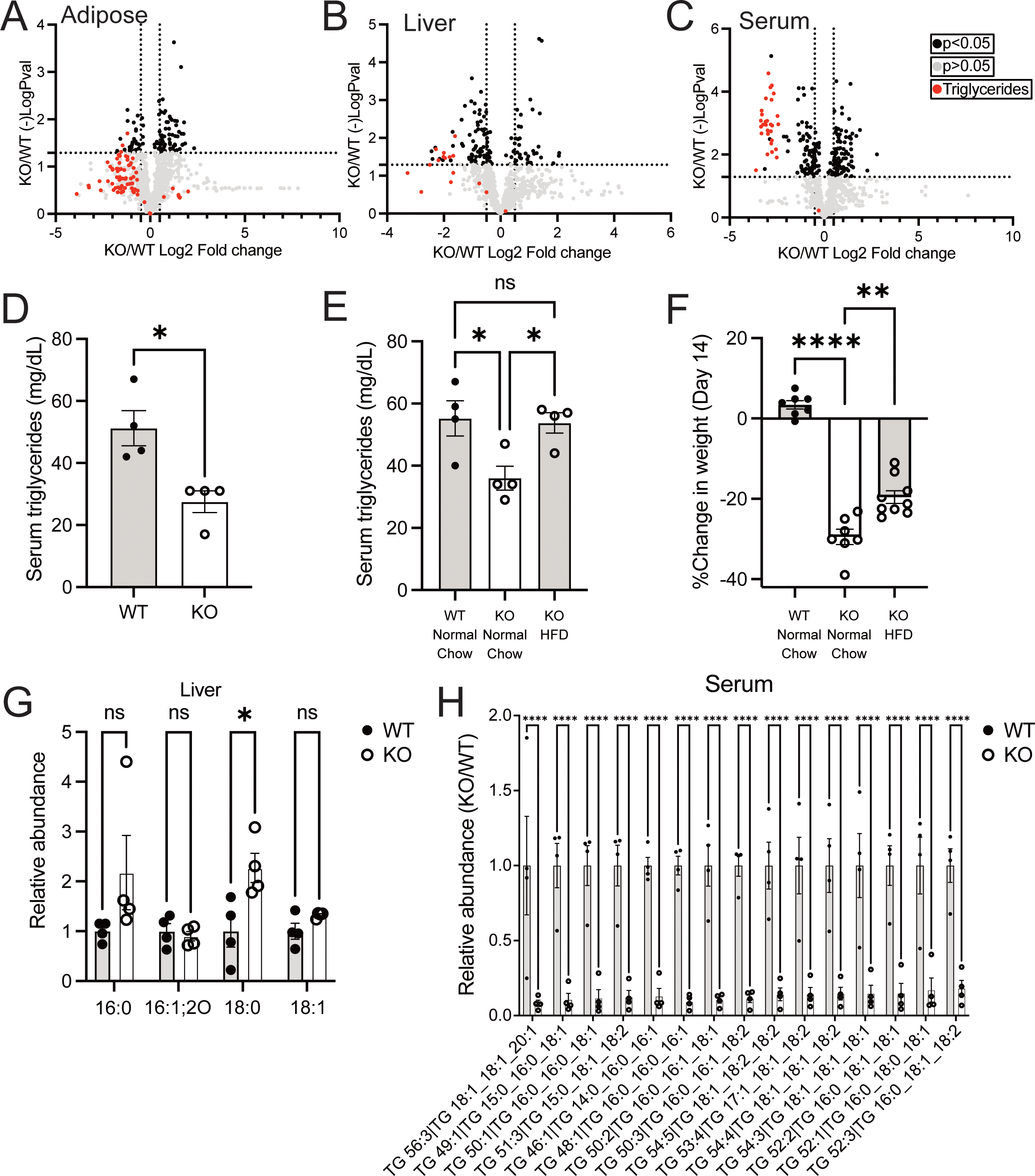
The abundance of triglycerides is lower following GSH depletion in mice. (A-C) Lipidomic analysis of (A) adipose tissue, (B) liver tissue, and (C) serum in KO (n=4) compared to WT (n=4) mice. Black data points = lipid species with a p <0.05 and log2 fold change > 2. Red data points = triglycerides. (D) Triglyceride levels in the serum of WT (n=4) and KO (n=4) mice (E) Serum triglyceride levels for WT (n=4) and KO (n=4) mice fed with either normal chow or a High Fat Diet (HFD) for 14 days post-treatment with tamoxifen. (F) Percent change in weight for WT (n=7) and KO mice fed with either normal chow (n= 7) or a high-fat diet (HFD; n=9) 14 days post-treatment with tamoxifen. (G-H) Relative abundance of select (G) fatty acids in the liver and (H) triglyceride species in the serum from WT (n=4) and KO (n=4) mice. 16:0 = palmitic acid; 16:1;2O= oxidized palmitoleic acid; 18:0 = stearic acid; 18:1 = oleic acid. Data are shown as mean ±SEM. An unpaired two-tailed t-test was used in (A-D), and a one-way ANOVA with subsequent Tukey’s multiple comparisons test was used in (E-H) to determine statistical significance. ns = not significant, * P value < 0.05, ** P value < 0.01, *** P value < 0.001, **** P value < 0.0001.

Next, we examined whether supplementing with a diet rich in lipids could reverse these reduced triglyceride and weight loss phenotypes. Supplying Gclc KO mice with a high-fat diet (HFD) rescued the decreased triglyceride levels in the serum **(Figure 3E)**. However, Gclc KO mice still lost substantial weight while on an HFD, although to a lesser extent than normal chow **(Figure 3F)**. These data suggest that maintenance of circulating triglycerides and sustaining body weight were potentially independent phenotypes in Gclc KO mice.

Loss of GSH synthesis resulted in a decreased expression of lipogenic enzymes, including SCD1. SCD1 catalyzes the desaturation of palmitic acid (16:0) into palmitoleic acid (16:1) and stearic acid (18:0) into oleic acid (18:1)^49^. Thus, we measured the abundance of these triglyceride species in the liver of Gclc KO mice. Levels of SCD1 substrates (i.e., palmitic acid and stearic acid) were elevated in the liver of Gclc KO mice **(Figure 3G)**. Further, the serum and liver from Gclc KO mice had decreased levels of triglycerides that contained palmitoleic acid and oleic acid **(Figure 3H and S3C)**. Together, these data demonstrate that ablation of GSH synthesis in mice results not only in reduced expression of lipogenic enzymes but also lower levels of their corresponding lipid species.

### Liver-specific GSH synthesis promotes lipid abundance

Whole-body ablation of GSH synthesis impacted lipid abundance *in vivo*. Synthesis of GSH and de novo lipids occurs in the liver. Whether liver-specific GSH synthesis was responsible for downregulated lipogenic gene expression and reduced serum triglycerides or whether these phenotypes were a byproduct of a liver-independent process in Gclc KO mice was unclear. Notably, Gclc KO mice rapidly lose weight and reach humane endpoints **(Figure 1E and 1F)**, making it difficult to extricate the other phenotypes potentially caused by the loss of GSH. To interrogate this, we utilized an AAV-TBG-Cre viral delivery system which produces liver-specific Cre expression and subsequent genetic deletion^50, 51^. *Gclc*^w/f^ or *Gclc*^w/w^ mice (Gclc WT) and *Gclc*^f/f^ mice (hereafter referred to as Gclc L-KO) were monitored following tail-vein injection with AAV-TBG-Cre **(Figure 4A)**. The liver from Gclc L-KO mice showed rapid depletion of *Gclc* mRNA and progressive loss of GCLC protein and GSH levels **(Figure 4B-4C)**. No differences in *Gclc* mRNA were observed in surrounding tissues **(Figure S4A-S4C)**, suggesting that the deletion of *Gclc* was localized to the liver. Unlike whole-body Gclc KO mice, liver-specific Gclc KO mice did not have any difference in survival, body weight, fat and lean mass, nor adipose tissue mass following *Gclc* deletion **(Figure 4D-4F and S4D-S4F)**. Gclc L-KO mice showed no histological signs of liver damage, although serum markers of liver damage were elevated compared to Gclc WT mice **(Figure 4G and S4G-S4H)**. These data demonstrate that deletion of the GCLC protein can be induced in the liver of adult animals. Importantly, these phenotypes contrast with the non-inducible liver-specific Gclc KO mouse strain (*Gclc*^f/f^ Albumin-Cre), which develops liver failure and dies shortly after birth^30^.

**Figure 4.**
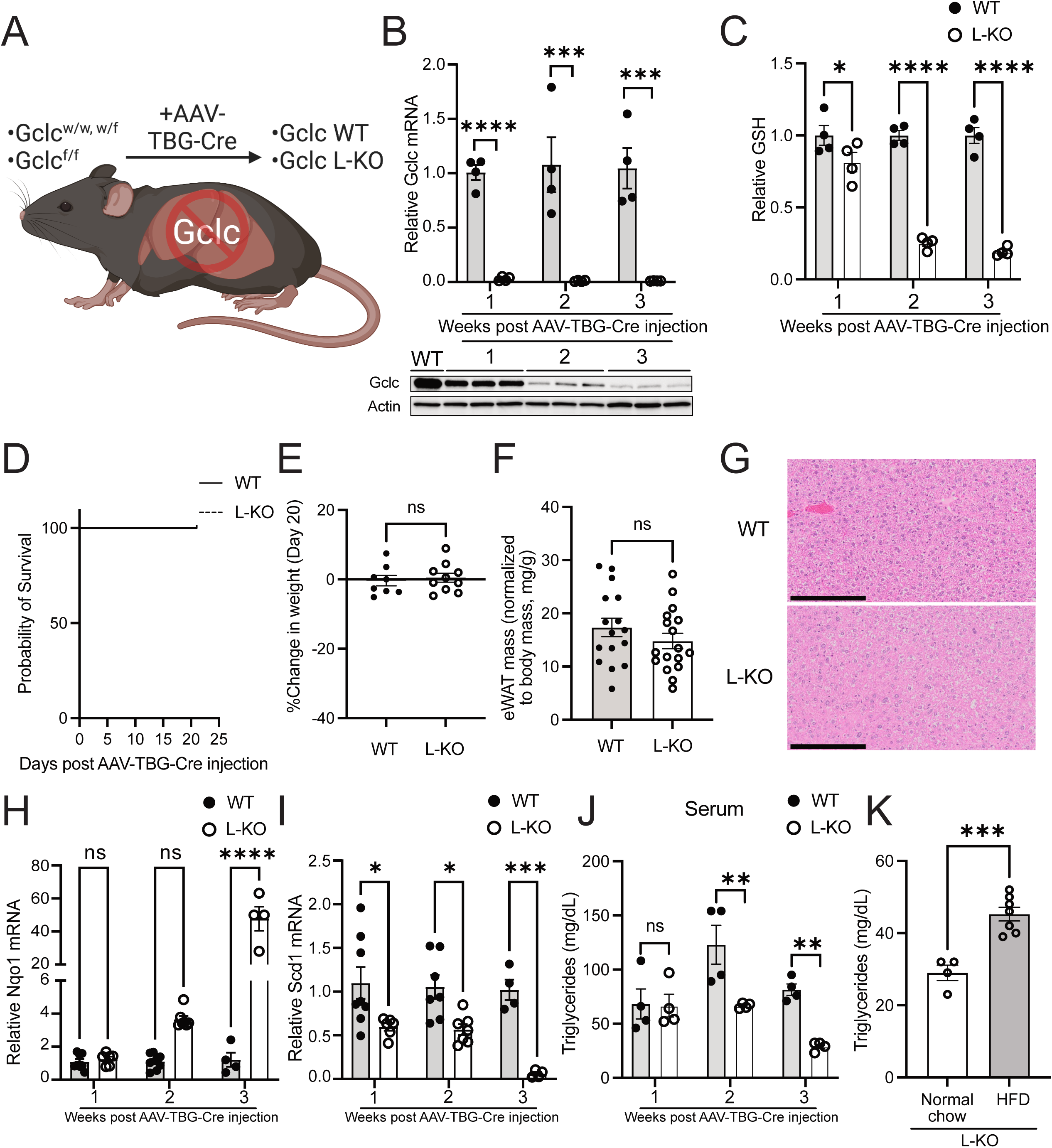
Liver-specific GCLC expression sustains lipid synthesis and represses NRF2 activation. (A) Schematic of inducible liver-specific *Gclc* deletion (L-KO). (B) Relative abundance of *Gclc* mRNA (top) and GCLC protein (bottom) in the liver of WT (n=4-8) and L-KO (n=4-7) mice following treatment with AAV-TBG-Cre. Expression levels of mRNA were normalized to the expression of the reference gene Rps9. (C) Relative abundance of GSH in liver tissue of WT (n=4) and L-KO (n=4) mice following treatment with AAV-TBG-Cre. (D) Percent survival of WT (n=6) and L-KO (n=6) mice following treatment with AAV-TBG-Cre. (E) Percent change in body weight of WT (n=8) mice and L-KO (n=10) mice at day 20 following treatment with AAV-TBG-Cre. (F) Epididymal white adipose tissue (eWAT) mass normalized to body mass in WT (n=16) and L-KO (n=17) mice. (G) Representative H&E-stained histochemical images of the liver of female WT and L-KO mice three weeks following treatment with AAV-TBG-Cre. Scale bars = 200 µm. (H-I) Relative mRNA levels of (H) Nqo1 and (I) Scd1 in WT (n=4-8) and L-KO (n=4-7) following treatment with AAV-TBG-Cre. Expression levels were normalized to the expression of the reference gene Rps9. (J) Triglyceride levels in the serum of WT (n=4) and L-KO (n=4) mice following treatment with AAV-TBG-Cre. (K) Triglyceride levels in the serum L-KO mice fed normal chow (n=4) and an HFD (n=7) 3 weeks following treatment with AAV-TBG-Cre. Data are shown as mean ±SEM. A one-way ANOVA with subsequent Tukey’s multiple comparisons test was used for (B-C) and (H-J), and an unpaired two-tailed t-test was used for (E-F) and (K) to determine statistical significance. ns =not significant,* P value< 0.05, ** P value< 0.01, *** P value < 0.001, **** P value< 0.0001.

Next, we examined whether livers from liver-specific Gclc KO mice induced an antioxidant response and had alterations in lipids and lipid-generating enzymes. Liver tissue from Gclc L-KO mice showed increased expression of NRF2 target genes and decreased lipogenic factors over time **(Figure 4H-I, S4I-S4L, and Table S4)**. Metabolomics revealed decreased expression of acetyl-CoA, an essential precursor for lipogenesis, and downstream acetyl-CoA metabolites **(Figure S4M)**. Further, we observed a gradual reduction in serum triglycerides in liver-specific Gclc KO mice, which was reversed when mice were placed on an HFD **(Figure 4J-5K)**. These data demonstrate that GSH synthesis in the liver is responsible for maintaining lipogenic gene expression and sustaining triglyceride levels *in vivo*.

**Figure 5:**
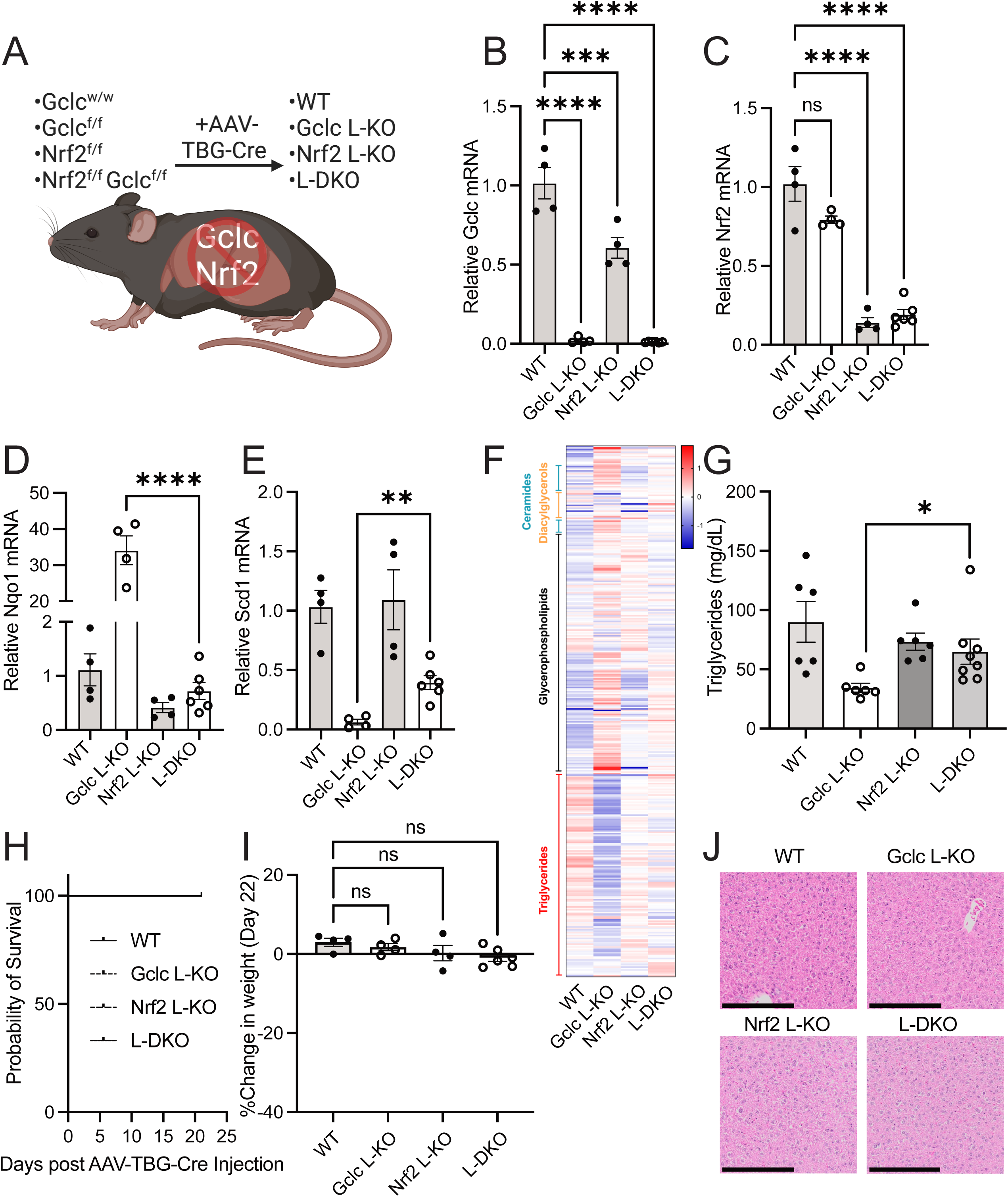
*Nrf2* deletion in the liver rescues triglyceride levels and lipid synthesis in liver- specific Gclc KO mice. (A) Schematic of inducible liver-specific *Gclc* deletion (Gclc L-KO), *Nrf2* deletion (Nrf2 L-KO), and *Gclc*-*Nrf2* deletion (L-DKO). (B-E) Relative expression of (B) *Gclc*, (C) *Nrf2*, (D) *Nqo1*, and (E) Scd1 mRNA in the liver of WT (n=4), Gclc L-KO (n=4), Nrf2 L-KO (n=4), and L-DKO (n=6) mice three weeks following treatment with AAV-TBG-Cre. Expression levels were normalized to the expression of the reference gene Rps9. (F) Lipidomic analysis of the serum of WT (n=4), Gclc L-KO (n=4), Nrf2 L-KO (n=4) and L-DKO (n=6) mice. Data are shown as normalized peak intensities. (G) Serum triglyceride levels for WT (n=6), Gclc L-KO (n=6), Nrf2 L-KO (n=6), and L-DKO (n=8) mice three weeks post-treatment with AAV-TBG-Cre. (H) Percent survival of WT, Gclc L-KO, Nrf2 L-KO & L-DKO following treatment with AAV-TBG-Cre. (I) Percent change in weight of WT (n=4), Gclc L-KO (n=4), Nrf2 L-KO (n=4), and L-DKO (n=6) mice three weeks following treatment with AAV-TBG-Cre. (J) Representative H&E-stained histochemical images of the liver of female WT, Gclc L-KO, Nrf2 L-KO, and L-DKO mice three weeks following treatment with AAV-TBG-Cre. Scale bars = 200 µm. Data are shown as mean ±SEM. A one-way ANOVA with subsequent Tukey’s multiple comparisons test was used for (B-C) and (I), and an unpaired two-tailed t-test was used for (D-E) and (G) to determine statistical significance. ns = not significant, * P value < 0.05, ** P value < 0.01, *** P value < 0.001, **** P value < 0.0001.

### GSH supports lipid abundance by repressing NRF2 activation

GSH is reported to contribute to lipid homeostasis in the body through several mechanisms^13, 52–56^. However, as shown above, our data suggest that the stabilization of NRF2 is associated with decreased expression of lipogenic enzymes and lower serum triglyceride levels. To test the involvement of NRF2, we bred the *Gclc*^f/f^ mouse strain with the *Nrf2*^f/f^ mouse strain^57, 58^ and induced a liver-specific double deletion with the AAV-TBG-Cre virus (hereafter referred to as L-DKO) **(Figure 5A)**. *Gclc* and *Nrf2* mRNA levels were decreased in L-DKO mice three weeks following injection of the AAV-TBG-Cre virus **(Figure 5B-5C)**, although the *Nrf2* gene was less efficiently deleted compared to the deletion of *Gclc*. Nonetheless, the deletion of *Nrf2* largely prevented the induction of NRF2 targets upon *Gclc* deletion in liver tissue **(Figure 5D and S5A)**. Further, the repression of lipogenic gene expression and the decreased serum triglyceride levels seen in Gclc L-KO mice were reversed in L-DKO mice **(Figure 5E-5G and S5B, Table S5)**. For specific lipogenic genes, the reversal of downregulated expression was not complete **(Figure 5E)**, suggesting the potential involvement of NRF2-independent pathways. Together, these results indicate that GSH supports lipid abundance by preventing NRF2 activation in the liver.

Activation of NRF2 prevents tissue damage under pathological conditions^59^. Since we observed an activation of NRF2 upon GSH depletion, we expected NRF2 to be essential for preventing liver damage. Surprisingly, L-DKO mice did not die or significantly lose weight **(Figure 5H-5I)**. Further, L-DKO mice did not show signs of liver failure **(Figure 5J and S5C)**, nor did they have increased levels of serum liver damage markers compared to Gclc L-KO mice **(Figure S5D)**. These results demonstrate that the expression of NRF2 in the liver is dispensable even in the absence of GSH. Further, they suggest that rather than buffering oxidative stress to prevent liver damage, GSH synthesis is crucial for maintaining low NRF2 levels and sustaining lipid production by the liver.

### Sustained depletion of liver GSH lowers fat stores

Liver-specific deletion of *Gclc* impaired the expression of lipogenic enzymes and lowered circulating triglyceride levels without causing liver failure. Adipose tissue, however, was not significantly depleted in liver-specific Gclc KO mice. Notably, these observations were found in a relatively short time after *Gclc* deletion (i.e., three weeks). We hypothesized that sustained repression of lipogenic programs would lower adipose stores over time. To test this, we monitored mice for ten weeks following the induction of *Gclc* deletion in the liver **(Figure 6A)**. Nearly all liver-specific Gclc KO mice survived and maintained weight over time **(Figure 6B-6C)**. Histological analysis revealed that for liver-specific Gclc KO mice, some livers appeared unharmed, while others showed signs of inflammation, cell death, and steatosis **(Figure 6D and S6A)**. However, there was no evidence of widespread damage across the liver tissues, and liver damage markers in the serum were not significantly elevated **(Figure S6B)**. Like the acute liver-specific *Gclc* deletion, serum triglycerides were depleted in mice with prolonged liver-specific *Gclc* deletion **(Figure 6E)**. Unlike the acute setting, however, prolonged ablation of GSH synthesis in the liver resulted in decreased abundance of white adipose depots **(Figure 6F and S6C)**. Interestingly, brown adipose tissue was not lower in liver-specific Gclc KO mice, potentially due to white and brown adipose tissues responding differently to redox changes^60^. These data demonstrate that GSH synthesis in the liver is required to maintain circulating essential for maintaining fat stores over time.

**Figure 6:**
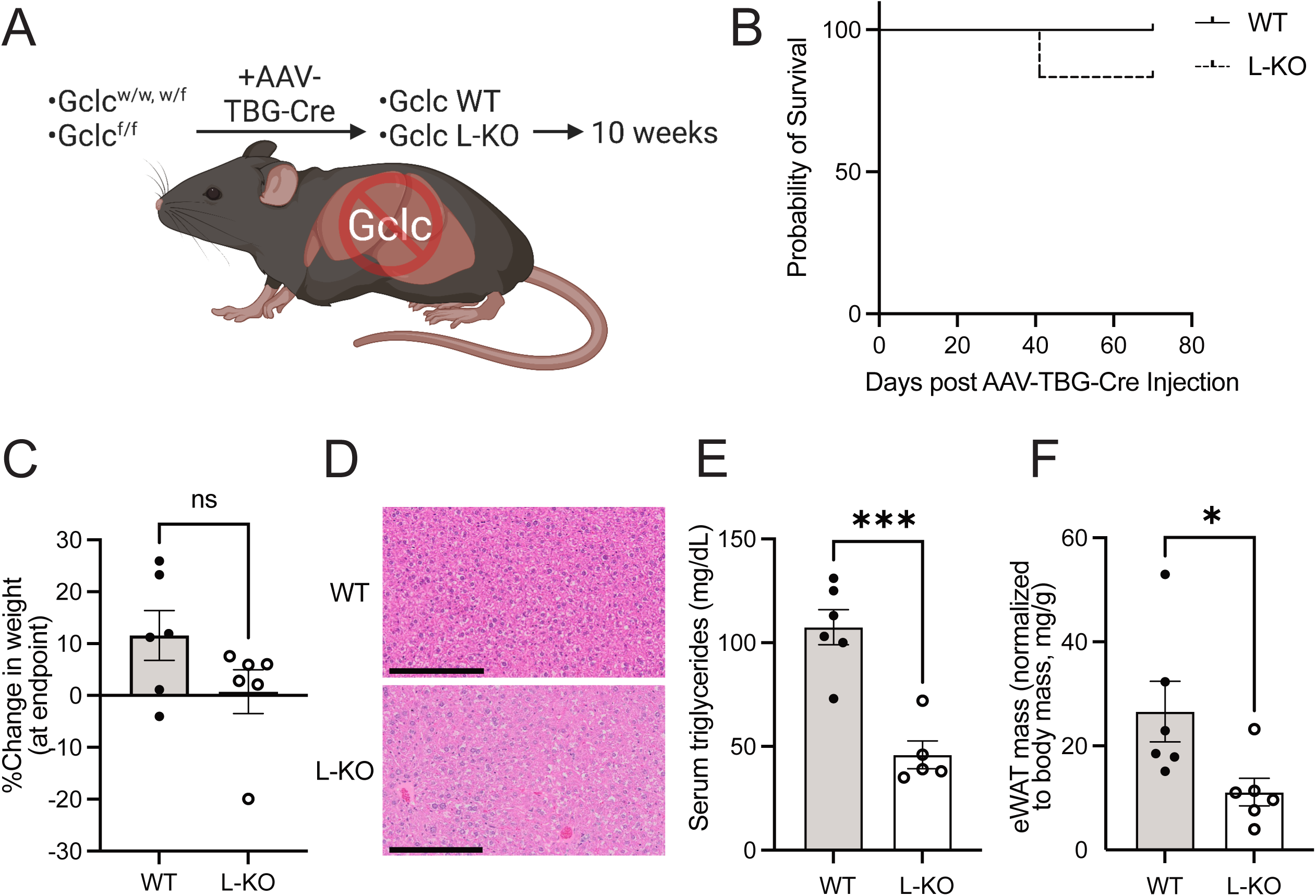
Prolonged *Gclc* deletion in the liver lowers adipose tissue mass. (A) Schematic for extended monitoring of WT and L-KO mice. (B) Percent survival for WT and L-KO mice following treatment with AAV-TBG-Cre. (C) Percent change in body weight at the endpoint of WT (n=6) and L-KO (n=6) mice. (D) Representative H&E-stained histochemical images of the liver from female WT and L-KO mice ten weeks following treatment with AAV-TBG-Cre. Scale bars = 200 µm. (E) Serum triglyceride concentration of WT (n=6) and L-KO (n=5) mice ten weeks following treatment with AAV-TBG-Cre. (F) Epididymal fat adipose tissue (eWAT) mass normalized to body mass from WT (n=6) and L-KO (n=6) mice ten weeks post-treatment with AAV-TBG-Cre. Data are shown as mean ±SEM. To determine statistical significance, an unpaired two-tailed t-test was used for (C) and (E-F). ns = not significant, * P value < 0.05, ** P value < 0.01, *** P value < 0.001, **** P value < 0.0001.

## DISCUSSION

GSH has been linked to nearly every cellular process in the body^61, 62^. However, its function *in vivo* remains poorly understood. To address this knowledge gap, we have developed a series of genetic models that permit spatiotemporal control over the expression of GCLC, the rate-limiting enzyme in GSH synthesis. We find that the loss of GCLC across all tissue triglycerides ands of adult animals results in rapid weight loss and death. However, this does not cause immediate damage to multiple tissues, including the liver. This is unexpected since we find the liver to have higher levels of GSH than other tissues. Previously, a liver-specific deletion of *Gclc* was shown to cause steatohepatitis and liver failure^30^. One difference between their model and the one reported here is that deletion of *Gclc* is induced in hepatocytes of adult mice (which are slowly proliferating), compared to theirs, where deletion of *Gclc* occurred in hepatocytes of growing animals (which are rapidly proliferating). The requirement of a protein pre- or post-organ and -animal development is an important distinction that is not always considered in studies involving tissue-specific knockout mouse models but potentially requires additional investigation.

A central question we ask here is, if GSH is not immediately required for liver survival, then why is it produced at such high levels? The clearest hypothesis is that GSH synthesis is required for liver function. Indeed, the liver has been reported to need a maximal synthesis of GSH for detoxifying xenobiotics, such as acetaminophen^63, 64^. Yet mammals are not constantly exposed to xenobiotics. Thus, we hypothesized that GSH synthesis plays additional detoxification-independent roles in liver function. The liver is known to be responsible for synthesizing new lipids. Also, lowering GSH levels in animals has previously been shown to prevent obesity-related phenotypes in mice, such as weight gain under a high-fat diet^65, 66^. In keeping with this observation, here we unveil a crucial role of GSH in lipid metabolism, as the synthesis of GSH, specifically in the liver, is required for sustaining physiological circulating triglyceride levels (an important energy source) and adipose tissue homeostasis under standard dietary regimen. This is a significant phenotype, as a similar reduction in circulating triglycerides was not observed with a liver-specific loss of critical lipogenic enzymes, such as fatty acid synthase (FASN)^67^. Together, our findings indicate that GSH synthesis in the liver is a crucial contributor to lipid abundance *in vivo*.

Mechanistically, we show that, upon GSH depletion, the liver tissue induces the expression of target genes associated with the antioxidant transcription factor NRF2. Since previous studies have linked the activation of Nrf2 with the repression of lipogenic gene expression^68^, we reasoned that GSH depletion causes oxidation and consequent inactivation of KEAP1 (a repressor of NRF2), thus leading to increased antioxidant and decreased lipogenic gene expression. However, KEAP1 regulates multiple pathways beyond NRF2^69, 70^, and the contributions of NRF2 to lipid homeostasis have been reported to be pleiotropic and context-dependent^71^. Chronic GSH deficiency by *Gclm* deletion in mice has also been associated with metabolic alterations and the activation of the stress-responsive factor AMPK^53, 72^. Further, AMPK has been reported to be an upstream activator of Nrf2^73^. Nonetheless, we show that following GSH depletion, the repression of lipogenic gene expression and serum triglyceride levels is dependent on NRF2 activation. Further investigations are required to elucidate why and how NRF2 activation blocks lipid production. One possible scenario is the metabolic cofactor NADPH, which is known to be required not only for lipid synthesis but also for the function of antioxidant reductases, such as glutathione reductase (GSR) and thioredoxin reductase (TXNRD)^74^. Notably, the deletion of lipogenic enzymes in the liver, such as acetyl-CoA carboxylase 1/2 (ACC1/2), has been reported to liberate NADPH for antioxidant processes, including GSH regeneration^75^.

In many scenarios, activation of NRF2 has been reported to be a stress response that is required to prevent the accumulation of oxidative stress and subsequent tissue damage. Surprisingly, we demonstrate here that, in an acute setting, mice with combined liver-specific deletion of *Gclc* and *Nrf2* do not lose weight or show poor health or liver failure. This suggests that the antioxidant response provided by NRF2 is not required for liver survival in the absence of GSH. Instead, our findings demonstrate a novel role for GSH in the liver, where under resting conditions, GSH maintains low NRF2 activity to permit ample expression of lipogenic enzymes, promotes the production of triglycerides, and sustains lipid abundance *in vivo*.

## METHODS

### EXPERIMENTAL MODEL AND SUBJECT DETAILS

#### Animal Studies

All animal studies were performed according to protocols approved by the University Committee on Animal Resources at the University of Rochester Medical Center. *Gclc*^f/f^ mice were generated as described^30^ and crossed with the Rosa26-CreERT2 mouse strain (Jackson Labs, #008463) or *Nrf2*^f/f^ mouse strain (Jackson Labs, #025433). All animals were aged for at least 12 weeks before being used in their respective experiments. Tamoxifen (Sigma Aldrich, T5648) was administered by intraperitoneal injection at 160 mg/kg once daily for five days. For liver-specific genetic deletions, mice were injected via the tail vein with 2.5×10^11^ GC of AAV-TBG-Cre (Addgene, 107787-AAV8) in PBS. For high-fat diet experiments, mice were fed a 60 kcal% fat high-fat diet (Research Diets, D12451i) following either tamoxifen or AAV-TBG-Cre injection for 2-3 weeks. Control mice were fed normal rodent chow (LabDiet, 5053).

#### Immunoblot assays

Tissue samples were crushed on dry ice to obtain homogenous aliquots and then lysed in RIPA buffer (Thermo Scientific #89900) containing Halt protease & phosphatase inhibitor (Thermo Scientific #1861280). Extracted proteins were quantified using the Pierce BCA Protein Assay Kit (Thermo Scientific #23225). 75 µg of protein lysates were heated for 10 mins at 95°C in Laemmli 6X SDS sample buffer (Boston BioProducts, # BP-111R) with 5% 2-mercaptoethanol (VWR Life Science #M131-100ml) and then ran on 4-20% Criterion TGX pre-cast gels (Bio-Rad #5671093). Separated proteins were transferred onto Immobilon-P Transfer membranes (MilliporeSigma #IPVH00010), blocked for an hour using 5% milk in TBST, and stained overnight with targeting primary antibodies in 5% milk TBST. Stained membranes were then washed with TBST and stained with corresponding secondary antibodies in 5% milk in TBST for one hour. Antibody-stained protein signal was amplified and visualized using SuperSignal™ West Pico PLUS chemiluminescent substrate (Thermo Fisher #34578) and imaged with a ChemiDoc MP Imaging system (Bio-Rad). The antibodies used for the immunoblot assays were: GCLC (Santa Cruz Biotech, #sc-390811), NQO1 (Sigma Prestige Antibodies, HPA007308), ACTIN (Sigma, A1978).

#### RNA analysis

Tissue samples were crushed on dry ice and homogenized using a bead Mill (VWR). mRNA was then isolated from the tissues using E.Z.N.A. total RNA Kit I (Omega Bio-Tek, R6834-02). For gene expression analysis, 1 µg of RNA was used for cDNA synthesis using qScript cDNA Synthesis Kit (Quanta Bio,#66196756). The expression of target genes was analyzed via Quantitative real-time (RT) PCR with a QuantaStudio 5 qPCR machine (Applied Biosystems, Thermo Fisher Scientific).

#### RNA-seq analysis

RNA-seq was performed on tissues using Genomics Research Center (GRC) at URMC and the Bauer Core Facility at Harvard University. Differential expression and GSEA analyses were performed as previously described^76, 77^.

#### Serum analysis

Mice were anesthetized with isoflurane, after which blood was collected via the retro-orbital venous sinus into BD microtainer tubes (BD #365967). Serum was isolated from the blood by centrifuging blood samples at 10000 xg for 5mins. Analysis of liver damage biomarkers and serum triglycerides was carried out by VRL Animal Health Diagnostics.

#### DEXA analysis

The fat and lean mass of mice were assessed with a dual-energy X-ray absorptiometry (DEXA) scanner (Lunar PIXImus2). Mice were initially anesthetized with isoflurane and then placed in a stretched-out prone position on the scanner bed, ensuring the tail and limbs do not touch the body. On the scanner bed, isoflurane was administered to maintain sedation. The lean and fat mass of the whole mouse body was measured via exposure to the X-ray from the scanner.

#### Tissue staining

Five-micron formalin-fixed, paraffin-embedded tissue sections were used for hematoxylin and eosin (H&E) staining and immunofluorescence analyses. The tissues were dewaxed and rehydrated through a series of xylene and ethanol changes. For antibody staining, antigen retrieval was performed on the slides by incubating them in a steamer for 40 mins in citrate buffer (Vector Labs, Cat# H-3300-250). The slides were washed in water and then blocked using 5% goat serum in PBS for 1 hr at room temperature prior to adding primary antibodies diluted in the blocking buffer. Primary antibody (NQO1; Abcam Cat# ab196196) incubation was carried out overnight at 4° C. For immunofluorescence, the second-to-last wash included 1 µg/ml of 4’, 6’-diamidino-2-phenylindole dihydrochloride (DAPI; Sigma, Cat# D9542-10mg) in PBS to stain cell nuclei. Immunofluorescence tissues were mounted with ProLong Gold (Life Technologies, Cat# P36934), while H&Es were mounted with Permount mounting media (Fisher Scientific, Cat# SP15500) and coverslipped for imaging. Immunofluorescence-stained slides were imaged using a CyteFinder II from Rarecyte. H&Es were imaged using an Olympus VS120 virtual slide microscope and Visiopharm image analysis system.

#### Preparation of NEM-derivatized cysteine and GSH internal standards

The N-ethylmaleamide (NEM) derivatized isotope labeled [^13^C_3_, ^15^N]-cysteine-NEM and [^13^C_2_,^15^N]-GSH-NEM were prepared by derivatizing the [^13^C_3_, 15N]-cysteine and [^13^C_2_,^15^N]-GSH standards with 50 mM NEM in 10 mM ammonium formate (pH = 7.0) at room temperature (30 min) as previously described^78^. [^13^C_4_, ^15^N_2_]-GSSG was prepared from the oxidation of [^13^C_2_,^15^N]-GSH as described^79^.

#### Non-targeted lipidomics – sample preparation

The liver, brain, kidney, lung, and adipose tissue were homogenized with a pre-chilled BioPulverizer (59012MS, BioSpec) and then placed on dry ice. The chloroform:methanol extraction solvent (v:v = 1:1) was added to the homogenate to meet 50 mg/mL. The samples were then sonicated in ice-cold water using Biorupter^TM^ UCD-200 sonicator for 5 min (30 s sonication and 30 s rest cycle; high voltage mode). The lipid extracts were cleared by centrifugation (17,000g, 20°C, 10 min), and the lipids in the supernatant were analyzed by LC-MS.

For serum samples, 75 µL of chloroform:methanol extraction solvent (v:v=1:2) was added to 20 µL of mouse serum, with the exception of the serum from WT, Gclc L-KO, Nrf2 L-KO, and L-DKO mouse serum, for which 20 µL of mouse serum was combined with 180 µL of chloroform:methanol extraction solvent (v:v=1:2) containing internal standards at the final concentrations: 5nM D7-Sphinganine (Avanti Polar Lipids Inc., Cat# 860658), 12.5nM D3-Deoxysphinganine (Avanti Polar Lipids Inc., Cat# 860474), and SPLASH LIPIDOMIX (1:1000, Avanti Polar Lipids Inc., Cat# 330707). After sonicating (1400 rpm, 20°C, 5 min), the extracts were cleared by centrifugation (17,000g, 20°C, 10 min), and the lipids in the supernatant were analyzed by LC-MS.

#### Non-targeted lipidomics – Instrumental condition and data analysis

The HPLC conditions were identical to the previous study^80^. In brief, chromatographic separation was conducted on a Brownlee SPP C18 column (2.1 mm × 75 mm, 2.7 μm particle size, Perkin Elmer, Waltham, MA) using mobile phase A (100% H2O containing 0.1% formic acid and 1% of 1 M NH4OAc) and B (1:1 acetonitrile:isopropanol containing 0.1% formic acid and 1% of 1 M NH4OAc). The gradient was programmed as follows: 0–2 min 35% B, 2–8 min from 35 to 80% B, 8–22 min from 80 to 99% B, 22–36 min 99% B, and 36.1–40 min from 99 to 35% B. The flow rate was 0.400 mL/min.

For the mass spectrometry, the data-dependent MS^2^ scan conditions were applied in both positive and negative mode: the scan range was from m/z 250–1500, resolution was 60,000 for MS, and 30,000 for DDMS^2^ (top 10), and the AGC target was 3E^6^ for full MS and 1E^5^ for DDMS^2^, allowing ions to accumulate for up to 200 ms for MS and 50 ms for MS/MS. For MS/MS, the following settings are used: isolation window width 1.2 m/z with an offset of 0.5 m/z, stepped NCE at 10, 15, and 25 a.u., minimum AGC 5E^2^, and dynamic exclusion of previously sampled peaks for 8 s.

For the analysis of lipids in the serum from WT, Gclc L-KO, Nrf2 L-KO, and L-DKO mice, the methods were the same with the following exceptions: MS2 scan conditions were applied in positive mode, the scan range was from m/z 120-1000, and the resolution was 120,000 for MS. For the MS/MS scan the following conditions were used: NCE at 20, 30, and 40 a.u. Quality control (QC) samples were included to check the technical variability and were prepared by mixing an equal volume of lipid extract from each tissue or serum sample. QC samples were included in the analysis sequence every ten samples and monitored for changes in peak area, width, and retention time to determine the performance of the LC-MS/MS analysis. QC samples were subsequently used to align the analytical batches.

The lipid peaks were identified, aligned, and exported using MS-DIAL ^81^. The data were further normalized to the median value of total lipid signals. Only lipids fully identified by MS^2^ spectra were included in the analysis.

#### Metabolomics – Sample preparation

The liver, brain, kidney, and lung tissues were homogenized with a pre-chilled BioPulverizer (59012MS, BioSpec) and then placed on dry ice. For the non-targeted metabolomics, the tissue metabolites were extracted in 80% MeOH at a final tissue concentration of 50 mg/mL for 24 hr at −80°C. For the quantification of sulfur metabolites in the tissue samples, the tissue metabolites were extracted and derivatized with NEM in ice-cold extraction solvent (80% MeOH: 20% H_2_O containing 25 mM NEM and 10 mM ammonium formate, pH=7.0) which includes stable isotope labeled internal standards (20 μM [^13^C_3_, ^15^N]-cysteine-NEM, 36.4 μM [^13^C_2_, ^15^N]-GSH-NEM, 10 μM [D_4_]-Cystine, 0.92 μM [^13^C_5_, ^15^N_2_]-GSSG, 20 μM [D_4_]-Hypotaurine, and 20 μM of [^13^C_2_]-Taurine at a final concentration of 50 mg/mL followed by incubation on 4°C for 24 hr.

For the global metabolite profiling in serum, the metabolites in 10 uL of serum were extracted by the 390 uL of pre-chilled 82% MeOH (−80°C) followed by 15 min incubation at −80°C. For the quantification of sulfur metabolites in serum, the metabolites in 20 uL of serum were extracted and derivatized by NEM in 80 uL of ice-cold extraction solvent (80% MeOH: 20% H2O containing 25 mM NEM and 10 mM ammonium formate, pH=7.0) which includes stable isotope labeled internal standards (1 μM [^13^C_3_, ^15^N]-cysteine-NEM, 0.1 μM [^13^C_2_, ^15^N]-GSH-NEM, 2 μM [D_4_]-Cystine, 4.6 μM [^13^C_5_, ^15^N_2_]-GSSG, 5 μM [D4]-Hypotaurine, and 40 μM [^13^C_2_]-Taurine followed by incubation at 4°C for 30 min.

After centrifugation (17,000 g, 20 min, 4C), all the supernatants were analyzed by LC-HRMS.

#### Metabolomics – Instrumental condition and data analysis

For the global metabolomics analysis, the previously established LC-MS conditions were applied^82^. For the chromatographic metabolite separation, the Vanquish UPLC systems were coupled to a Q Exactive HF (QE-HF) mass spectrometer equipped with HESI (Thermo Fisher Scientific, Waltham, MA). Samples were run on either a SeQuant ZIC-pHILIC LC column, 5 mm, 150 x 4.6mm (MilliporeSigma, Burlington, MA) with a SeQuant ZIC-pHILIC guard column, 20 x 4.6 mm (MilliporeSigma, Burlington, MA) or an Atlantis Premier BEH Z-HILIC VanGuard FIT column, 2.5µm, 2.1mm x 150mm (Waters, Milford, MA). For all samples, mobile phase A was 10mM (NH_4_)_2_CO_3_ and 0.05% NH_4_OH in H_2_O, while mobile phase B was 100% ACN. The column chamber temperature was set to 30°C. The mobile phase condition was set according to the gradient of 0-13min: 80% to 20% of mobile phase B, 13-15min: 20% of mobile phase B. The ESI ionization mode was positive and negative. For samples run on the ZIC-pHILIC column, the MS scan range (m/z) was set to 60-900. For those run on the BEH X-HILIC column, the MS scan range was set to 65-975. The mass resolution was 120,000, and the AGC target was 3 x 10^6^. The capillary voltage and capillary temperature were set to 3.5 KV and 320°C, respectively. 5 μL of the sample was loaded. The LC-MS metabolite peaks were manually identified and integrated by EL-Maven (Version 0.11.0) by matching with a previously established in-house library ^82^.

For the targeted sulfur metabolite quantification approach, the previously established LC-MS conditions were applied ^82^ with selected reaction monitoring (MRM) using an Ultimate 3000 UPLC system coupled to a Thermo Finnigan TSQ Quantum equipped with HESI (Thermo Fisher Scientific, Waltham, MA). As a stationary phase, an XBridge Amide Column 3.5 µm (2.1 × 100mm) (Waters, Milford, MA) was used. The mobile phase A was 97% water and 3% ACN (20 mM NH_4_Ac, 15 mM NH_4_OH, pH = 9.0), and the mobile phase B was 100% ACN. The column temperature was set to 40°C, and the gradient elution was at 0.35 mL/mL of flow rate: 0 to 3 min, linear gradient from 15% to 70% of Phase A; 3 to 12 min: linear gradient from 70% to 98% of Phase A; 12 to 15 min, sustaining 98% of Phase A. The MS acquisition operated in the positive or negative mode. The capillary temperature was 305 °C, and the vaporizer temperature was 200 °C. The sheath gas flow was 75, and the auxiliary gas flow was 10. The spray voltage was 3.7 kV. The MRM conditions (parent ion → fragment ion; collision energy) of metabolites were as follows. Positive mode: Cysteine-NEM (m/z 247 → m/z 158; 30); [^13^C_3_, ^15^N]-Cysteine-NEM (m/z 251 → m/z 158; 30); GSH–NEM m/z (m/z 433 → m/z 304; 15); [^13^C_2_,^15^N]-GSH-NEM (m/z 436 → 307m/z; 15); Cystine (m/z 241→ m/z 74; 30); [D_4_]- Cystine (m/z 245 → m/z 76; 30); GSSG (m/z 613 → m/z 355; 25), [^13^C_4_, ^15^N2]-GSSG (m/z 619 → m/z 361; 25); Hypotaurine (m/z 110 → m/z 92; 10); [D_4_]-Hypotaurine (m/z 114 → m/z 96; 10); Taurine (m/z 126 → m/z 108; 11); [^13^C_2_]-Taurine – (m/z 128 → m/z 110; 11). All peaks were manually integrated using Thermo Xcaliber Qual Browser. The quantification of metabolites was calculated by an isotope ratio-based approach according to published methods^83^.

#### Global protein abundance profiling by mass spectrometry

##### Sample preparation (A)

Flash frozen liver sections (∼25 mg) from WT or *Gclc*^-/-^ mice (3 biological replicates each per TMT sample) were thawed on ice and resuspended in DPBS, supplemented with protease (cOmplete™, EDTA-free protease inhibitor cocktail, Roche, #11873580001) and phosphatase (PhosSTOP™, Roche, #4906845001) inhibitor tablets. Tissue was homogenized by probe sonication (2 x 10 pulses, 40% power output), and particulate matter was removed by passing samples through 0.4 μm syringe filters. The proteome concentration of the tissue lysates was determined using the DC protein assay (Bio-Rad), normalized to 2 mg/mL, and then 100 μL of each sample was transferred to a LoBind Eppendorf tube containing 48 mg of urea. Samples were reduced with DTT (5 μL of 200 mM stock in H_2_O, 10 mM final concentration) and incubated at 65 °C for 15 minutes, then alkylated with iodoacetamide (5 μL of 400 mM stock in H_2_O, 20 mM final concentration) and shaken at 37 °C for 30 minutes in the dark. Ice-cold MeOH (600 μL), CHCl_3_ (200 μL), and H_2_O (500 μL) samples were vortexed and then centrifuged (10,000 g, 10 minutes, 4 °C) to precipitate proteins. The upper layer of supernatant was removed, and ice-cold MeOH (600 μL) was added to wash the protein disc. Samples were vortexed again, centrifuged (16,000 g, 10 minutes, 4°C) and then all supernatant was removed to leave a protein pellet. Samples were resuspended in 160 μL EPPS buffer (200 mM, pH 8.0) using a probe sonicator (1 x 10-15 pulses, power output 20%) and then digested with LysC (4 μL of 0.5 μg/μL per sample, resuspended in HPLC grade water, Wako-chemicals, Fujifilm #125-05061) for 2 h at 37 °C in a shaker incubator. Samples were then digested with trypsin (11 μL of 0.5 μg/μL per sample, resuspended in trypsin resuspension buffer containing 20 mM CaCl_2_; Promega, #V542A) overnight at 37 °C in a shaker incubator. After incubation with trypsin, the peptide concentration in samples was estimated using a Micro BCA^TM^ Protein Assay (Thermo Scientific, #23235), and a volume corresponding to 25 μg peptides was transferred to a new low-bind Eppendorf tube per sample. Sample volumes were normalized to 35 μL with EPPS buffer (200 mM, pH 8), diluted with HPLC grade CH_3_CN (9 μL), and then labeled (5 μL of 20 μg/μL per sample) with the corresponding TMTsixplex™ Isobaric Mass Tag (Themo Scientific, #90064B). Samples were incubated at room temperature for 1 h, vortexing intermittently, and then quenched by the addition of hydroxylamine (5 μL of 5% w/v in HPLC water per sample) and incubated for 15 minutes at room temperature. Samples were then acidified with formic acid (2.5 μL), and 2 µL of each sample was combined in a LoBind Eppendorf and dried using a Speedvac to perform a ratio check. Remaining samples were stored at −80 until after experimentally determining TMT channel intensities.

##### Sample preparation (B)

In an alternate protocol, the frozen liver tissue was directly lysed by probe sonication (2 x 10 pulse, 40% power output) in 4 M urea/DPBS and briefly cleared of debris by centrifugation (5000 g, 5 min, 4 °C). The proteome concentration of lysates was normalized to 2 mg/mL, and 100 μL was transferred to a LoBind Eppendorf tube containing 48 mg of urea. Samples were reduced with DTT and alkylated with iodoacetamide as described above, then diluted with DPBS (300 μL), and taken forward to LysC and trypsin digestion without precipitation. Digested samples were desalted using a C18 spin column (Pierce™ C18 Spin, #89873) according to manufacturer instructions prior to resuspension in EPPS buffer and CH_3_CN for TMT labeling as described above.

##### TMT ratio check

The combined and dried “ratio check” sample was re-dissolved in Buffer A (5% CH_3_CN, 95% water, 0.1% formic acid, 20 μL) and desalted using C18 stage tips prepared in-house using 3 x C18 discs (3M Empore) stacked in 200 μL pipette tips. Stage-tips were activated with MeOH (2 x 60 μL), washed with Buffer B (80% CH_3_CN, 20% water, 0.1% formic acid) (1 x 60 μL), and equilibrated with Buffer A (2 x 60 μL). The entire sample was then passed through the stagetip twice before being eluted into a new LoBind Eppendorf using 80 μL of 70% CH_3_CN/30% H_2_O/0.1% formic acid). The desalted sample was evaporated to dryness in a speedvac and then resuspended in 10 μL Buffer A and analyzed by mass-spectrometry using the following LC-MS gradient: 5% buffer B in buffer A from 0-15 min, 5%–15% buffer B from 15-17.5 min, 15%–35% buffer B from 17.5-92.5 min, 35%–95% buffer B from 92.5-95 min, 95% buffer B from 95-105 min, 95%–5% buffer B from 105-107 min, and 5% buffer B from 107-125 min; and standard MS3-based quantification described below. Ratios were determined from the average peak intensities corresponding to each channel. After experimentally determining TMT channel signal intensities, frozen samples were thawed, and a volume corresponding to 12.5 μg/sample was combined in a new LoBind Eppendorf tube and dried using a SpeedVac.

##### High pH fractionation

TMT labeled, combined, and dried samples were resuspended in 300 μL of Buffer A (5% v/v MeCN, 95% v/v H_2_O, 0.1% v/v formic acid) and fractionated by centrifugation using a peptide desalting column (Pierce™, #89852). Briefly, the storage solution was removed (5,000 g, 2 min), and columns were washed with CH3CN (2 x 300 μL, 5,000 g, 2 min), then equilibrated with buffer A (2 x 300 μL, 5,000 g, 2 min). Resuspended samples were loaded onto equilibrated spin columns (2,000 g, 2 min), passing the entire sample twice through the column. The column was then washed with buffer A (300 μL, 2,000 g, 2 min), and 10 mM aqueous NH4HCO3 containing 5% CH3CN (300 μL, 2,000 g, 2 min), before peptides were eluted from the column as 15 fractions using 300 μL buffer containing an increasing concentration of CH_3_CN in 10 mM NH4HCO3 (%CH_3_CN = 7.5, 10, 12.5, 15, 17.5, 20, 22.5, 25, 27.5, 30, 35, 40, 45, 50, 75) (2000 g, 2 min). Every 5^th^ fraction was combined into a new Eppendorf to make 5 final fractions that were dried using a SpeedVac vacuum concentrator. The resulting fractions were then re-suspended in buffer A (16 μL) and analyzed on an Orbitrap Fusion mass spectrometer.

##### Data processing

Protein abundance was calculated as a ratio of *Gclc*^-/-^ vs. WT samples for each peptide-spectra match by dividing each TMT reporter ion intensity by the average intensity for the channels corresponding to WT control (3 per TMT sample). Peptide-spectra matches were then grouped based on protein ID, excluding peptides with summed reporter ion intensities for the WT channels < 15,000, coefficient of variation for WT channels > 0.5, non-unique or non-tryptic peptide sequences. TMT reporter ion intensities were normalized to the median summed signal intensity across channels, and the data were filtered to retain proteins with at least 2 distinct peptides. The fold change in protein abundance in KO vs. WT mice was calculated for individual replicates, averaged, and converted to a log2 scale. Change in proteins abundance ≥ 1 log unit and p-value <0.05 were considered significant.

#### TMT liquid chromatography-mass-spectrometry (LC-MS) analysis

Samples were analyzed by liquid chromatography tandem mass spectrometry using an Orbitrap Fusion Tribrid Mass Spectrometer (Thermo Scientific) coupled to an UltiMate 3000 Series Rapid Separation LC system and autosampler (Thermo Scientific Dionex). The peptides were eluted onto an EASY-Spray HPLC column (Thermo ES902, ES903) using an Acclaim PepMap 100 (Thermo 164535) loading column and separated at a flow rate of 0.25 μL/min. Data were acquired using an MS3-based TMT method using the following scan parameters: scan sequence began with an MS1 master scan (Orbitrap analysis, resolution 120,000, 400−1700 m/z, RF lens 60%, automatic gain control [AGC] target 2E5, maximum injection time 50 ms, centroid mode) with dynamic exclusion enabled (repeat count 1, duration 15 s). The top ten precursors were then selected for MS2/MS3 analysis. MS2 analysis consisted of quadrupole isolation (isolation window 0.7) of precursor ion followed by collision-induced dissociation (CID) in the ion trap (AGC 1.8E4, normalized collision energy 35%, maximum injection time 120 ms). Following the acquisition of each MS2 spectrum, synchronous precursor selection (SPS) enabled the selection of up to 10 MS2 fragment ions for MS3 analysis. MS3 precursors were fragmented by HCD and analyzed using the Orbitrap (collision energy 55%, AGC 1.5E5, maximum injection time 120 ms, resolution was 50,000). For MS3 analysis, we used charge state–dependent isolation windows. For charge state z = 2, the MS isolation window was set at 1.2; for z = 3-6, the MS isolation window was set at 0.7. The RAW files were uploaded to Integrated Proteomics Pipeline (IP2) and searched using the ProLuCID algorithm (publicly available at http://fields.scripps.edu/yates/wp/?page_id=821) using a reverse concatenated, non-redundant version of the Mouse UniProt database (release 2017). Cysteine residues were searched with a static modification for carboxyamidomethylation (+57.02146 Da), and N-termini and lysine residues were searched with a static modification corresponding to the TMT tag (+229.1629 Da). Methionine residues were searched with a differential modification for oxidation (+15.9949 Da) and a maximum of 4 differential modifications were allowed per peptide. Peptides were required to be at least 5 amino acids long and fully tryptic. ProLuCID data was filtered through DTASelect (version 2.0) to achieve a peptide false-positive rate below 1%. The MS3-based peptide quantification was performed with reporter ion mass tolerance set to 30 ppm with Integrated Proteomics Pipeline (IP2).

## STATISTICAL ANALYSIS

All statistical analysis was completed using either R or GraphPad Prism 9

## DATA AVAILABILITY

Data supporting these findings are included within article and its supplementary material. Raw data supporting the findings reported in this study are available from the corresponding author upon request

## ACKNOWLEDGMENTS

We thank Jonathan Coloff and Samuel McBrayer for their feedback and discussions. We would also like to thank the Genomics Research Center (GRC), the Histology, Biochemistry, and Molecular Imaging (HBMI) Core at the Center for Musculoskeletal Research (CMSR), and the Center for Advanced Research Technologies (CART) at URMC, the Bauer Core Facility at Harvard University, and Proteomics/Metabolomics Core at Moffitt Cancer Center, which is funded in part by Moffitt’s Cancer Center Support Grant (P30CA076292). We also thank Joan Brugge and the Ludwig Cancer Center at Harvard Medical School for their support. This work was supported by the American Association for Cancer Research and Breast Cancer Research Foundation (20-20-26-HARR) (I.S.H.), Breast Cancer Coalition of Rochester (I.S.H.), NIH grants R01CA269813 (I.S.H.), R37CA230042 (G.M.D), R24AA022057 (V.V.), R01AA028859 (Y.C.), AI150698 (J.M.), and R01AR078000 (R.T.D), and a Sir Henry Wellcome Postdoctoral Fellowship (M.E.K.). Schematics were created with BioRender.com.

## AUTHOR CONTRIBUTIONS

G.A. and I.S.H. initiated the study, conceived the project, designed experiments, interpreted results, and wrote the manuscript. G.A. performed the experiments with assistance from E.T.T. for animal breeding and experiments. N.P.W., Y.P.K., and G.M.D. performed metabolomic and lipidomic analyses. M.E.K. and B.F.C. performed proteomic analyses. N.G. performed immunofluorescent experiments. R.D, K.R., F.H., M.Z., L. S-S., T.Q.S., K.T., F.A., Z.R.S. assisted with animal necropsies. D.A-V., A.F.H., A.R.H., T.N.O., R.T.D., S.W., A.R., R.T.B., J.C., G.K.G., assisted with animal experiments and histological analyses. L.M.S. performed bioinformatic analyses. K.R. assisted with RNA analysis. H.C. and Z.S. assisted with immunoblot experiments. C.C. assisted with DEXA analyses. Y.C., V.V., D.B., X.L.S, and J.M. provided expert comments and reagents.

---

**Figure S1.**
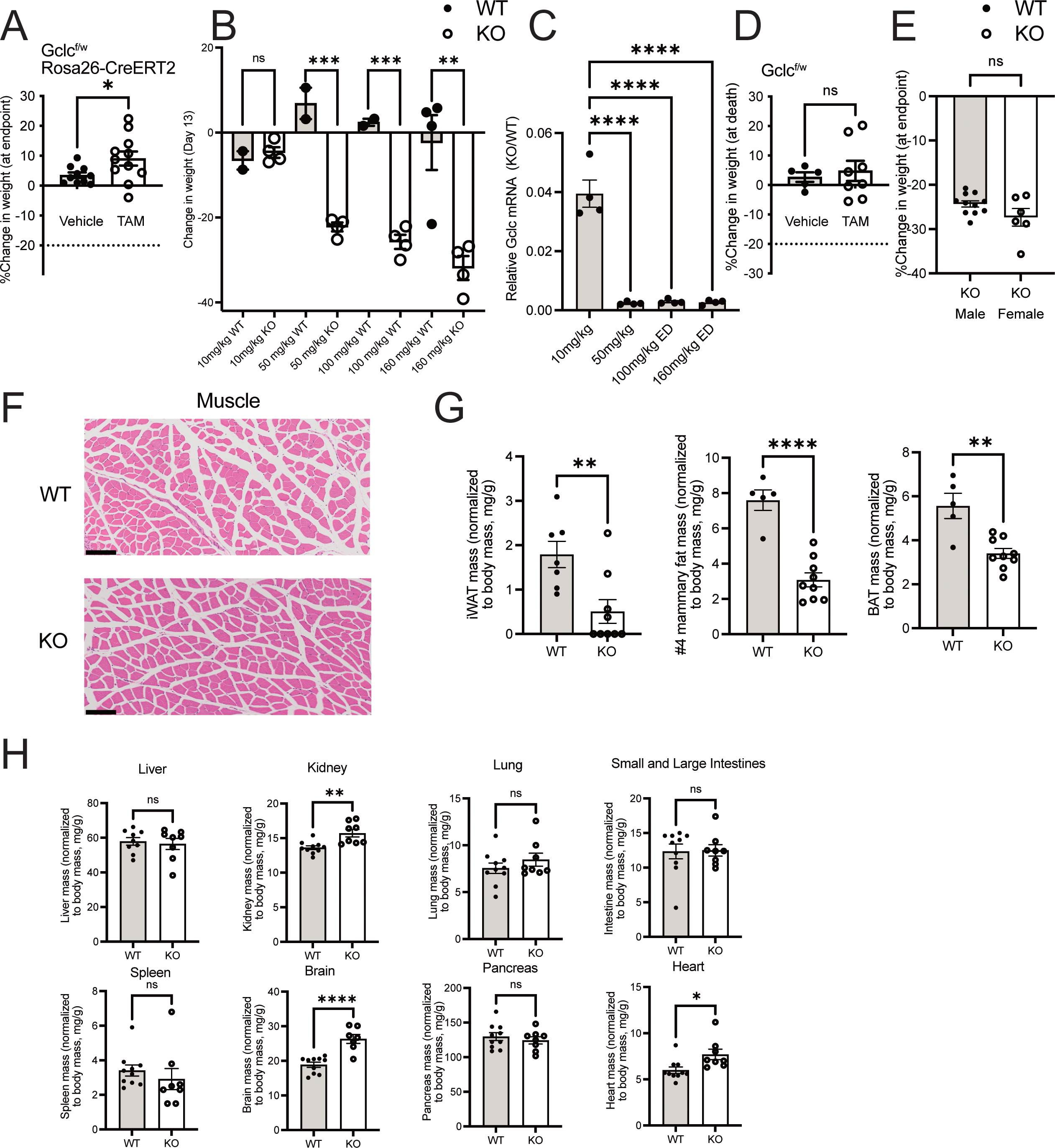
GSH depletion disproportionally affects adipose tissue. Related to Figure 1. (A) Percent change in body weight in *Gclc*^f/w^ Rosa26-CreERT2 mice treated with either vehicle (corn oil) (n=10) or tamoxifen (n=11). The dotted line indicates a 20% cutoff weight change for humane endpoints. (B) Percent change in body weight at endpoint between WT (n=2-4) and KO (n=4) mice treated with varying tamoxifen doses (10, 50, 100, and 160 mg/kg body weight). (C) Relative expression of *Gclc* mRNA in the liver from the KO (n=4) compared to WT (n=4) mice treated with different doses of tamoxifen (10, 50, 100, and 160 mg/kg body weight). (D) Percent change in body weight at the endpoint in *Gclc*^f/w^ mice treated with vehicle (corn oil) (n=4) or tamoxifen (n=8). (E) Percent change in body weight at the endpoint in male (n= 11) and female KO (n=5) mice. (F) Representative H&E-stained histochemical images of the muscle tissue from WT (top) and KO (bottom) mice (scale bars = 200 µm) (G) Inguinal fat, #4 mammary fat, and brown adipose tissue (BAT) mass normalized to body mass from WT (n=5-7) mice and KO (n=9) mice. (J) Indicated tissue mass normalized to body mass from WT (n = 9-10) mice and KO (n=7-8) mice. Data are shown as mean ±SEM. A one-way ANOVA with subsequent Tukey’s multiple comparisons test was used for (B-C) and (I), and an unpaired two-tailed t-test was used for (A), (D-E), and (G-H) to determine statistical significance. ns = not significant, * P value < 0.05, ** P value < 0.01, *** P value < 0.001, **** P value < 0.0001.

**Figure S2.**
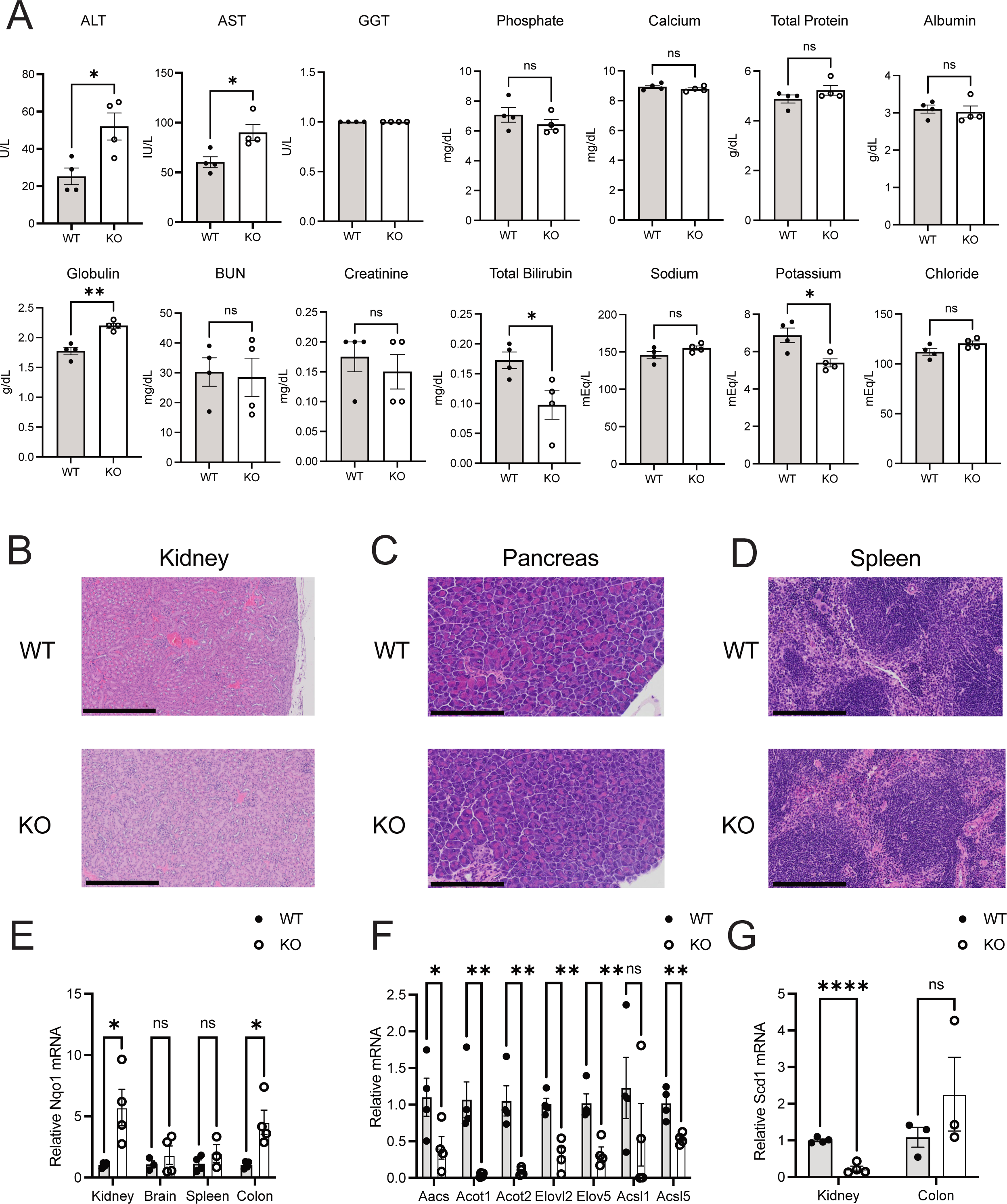
Ablation of GSH synthesis in animals does not cause damage to multiple tissues. Related to Figure 2. (A) Biochemical analysis of annotated serum biomarkers from WT (n=4) and KO (n=4) mice. (B-D) Representative H&E-stained histochemical images of the (B) kidney, (C) pancreas, and (D) spleen from WT mice (top) and KO (bottom) mice. Scale bars = (B) 500 µm, (C-D) 200 µm. (E) Relative expression of Nqo1 mRNA in the kidney, brain, spleen, and colon from WT (n=4) and KO (n=4) mice. Expression levels were normalized to the expression of the reference gene Rps9. (F) Relative mRNA expression of annotated lipogenic genes in the liver of WT (n=4) and KO (n=4) mice. Expression levels were normalized to the expression of the reference gene Rps9. (G) Relative expression of Scd1 mRNA in the kidney and colon from Gclc WT (n=4) mice and Gclc KO (n=4) mice. Expression levels were normalized to the expression of the reference gene Rps9. Data are shown as mean ±SEM. A one-way ANOVA with subsequent Tukey’s multiple comparisons test was used for (E-G), and an unpaired two-tailed t-test was used for (A) to determine statistical significance. ns = not significant, * P value < 0.05, ** P value < 0.01, *** P value < 0.001, **** P value < 0.0001.

**Figure S3.**
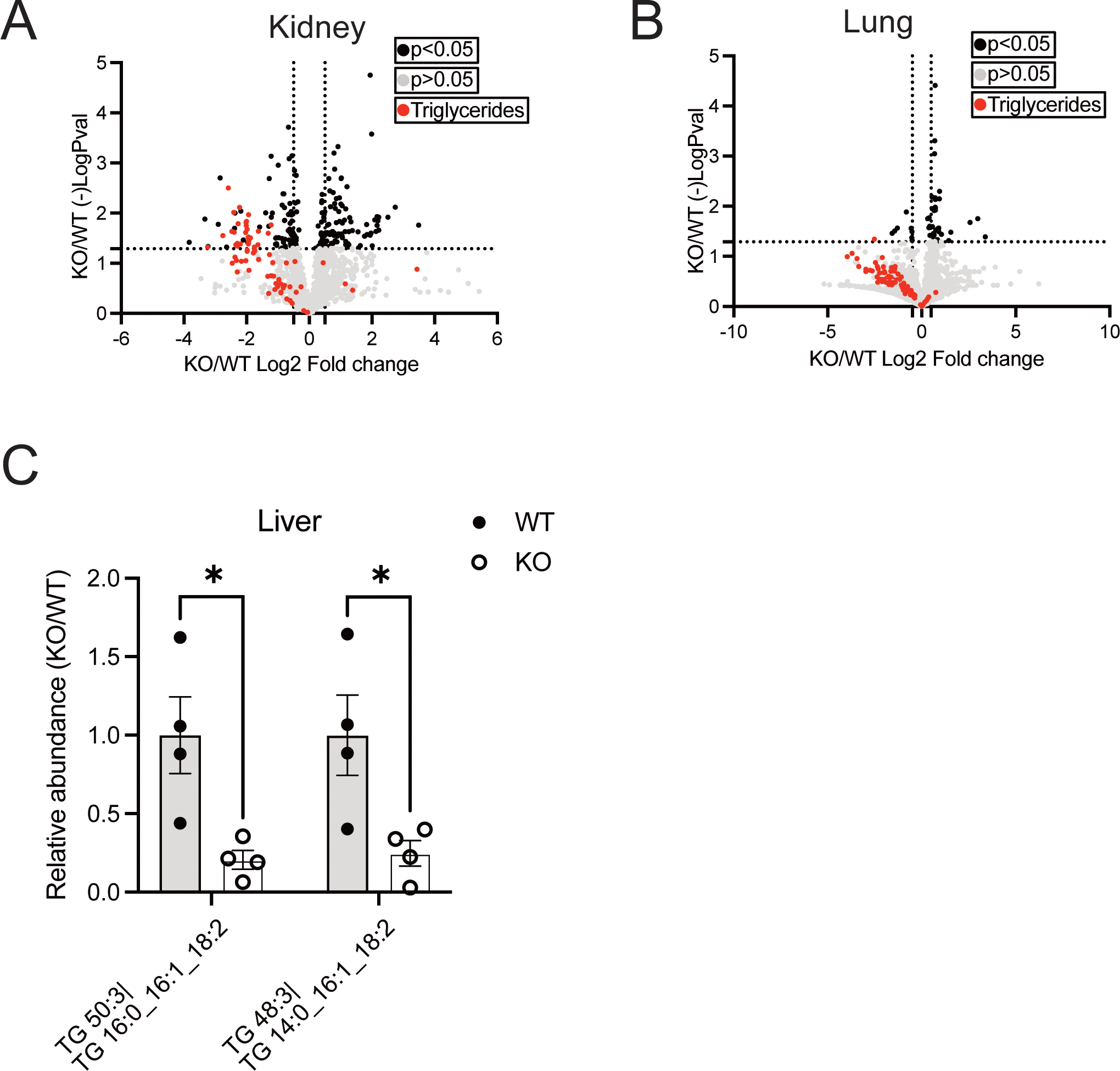
Blocking GSH synthesis lowers the abundance of triglycerides in tissues. Related to Figure 3. (A-B) Fold change of lipid species from the kidney (A) and lung (B) of Gclc WT and Gclc KO mice. Black data points = lipid species with a p <0.05 and log2 fold change > 2. Red data points = triglycerides. (C) Relative abundance of annotated triglyceride species in the liver of WT (n=4) and KO (n=4) mice. Data are shown as mean ±SEM. A one-way ANOVA with subsequent Tukey’s multiple comparisons test was used for (C), and an unpaired two-tailed t-test was used for (A-B) to determine statistical significance. ns = not significant, * P value < 0.05, ** P value < 0.01, *** P value < 0.001, **** P value < 0.0001.

**Figure S4.**
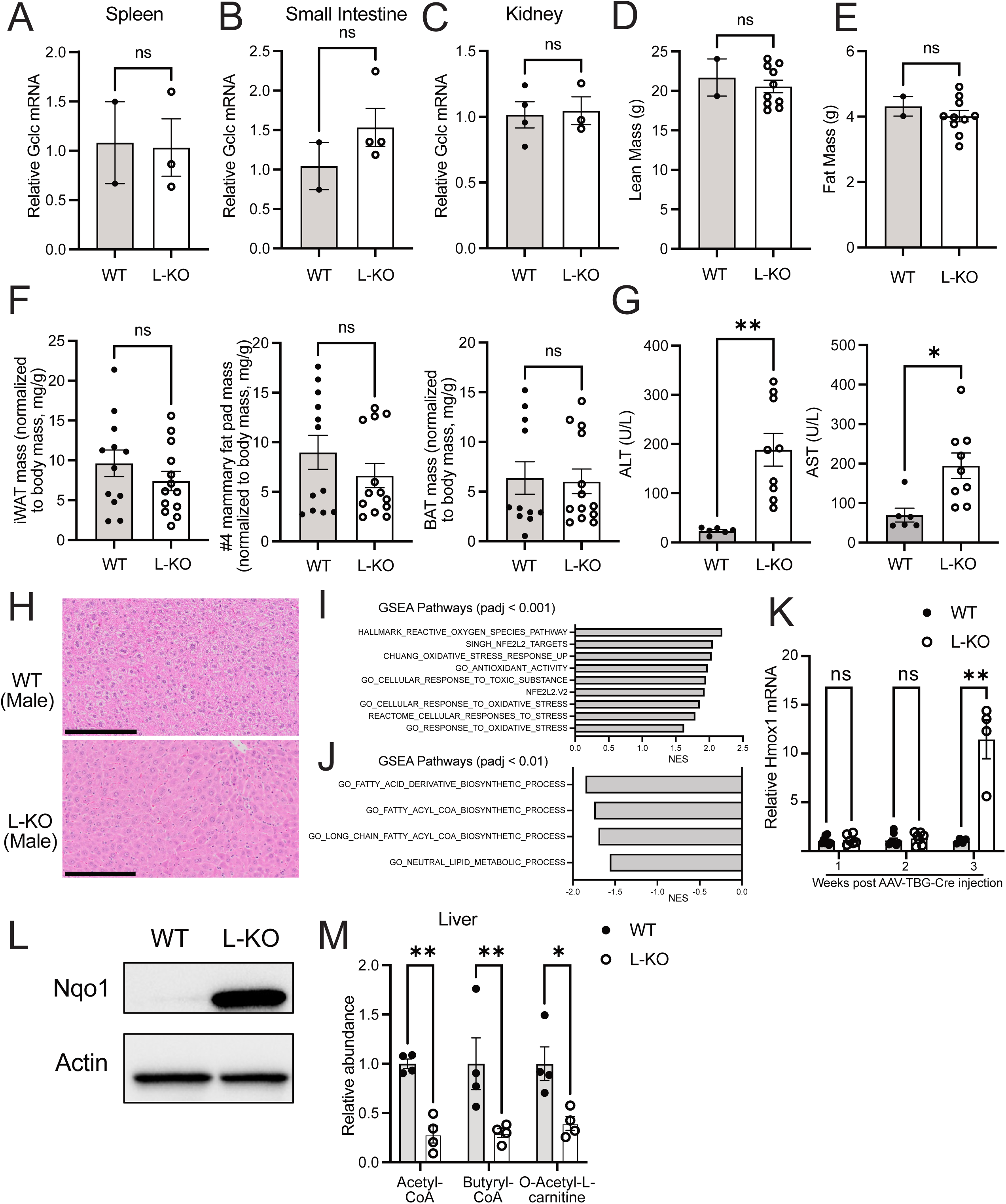
GSH produced in the liver promotes triglyceride abundance and suppresses NRF2 activity. Related to Figure 4. (A-C) Relative expression of *Gclc* mRNA in the (A) spleen, (B) small intestine, and (C) kidney from WT (n=2-4) and L-KO (n=3-4) mice. Expression levels were normalized to the expression of the reference gene Rps9. (D-E) Dexa analysis of (D) fat mass and (E) lean mass from WT (n=2) mice and L-KO (n=9-10) (F) Tissue mass normalized to body mass of inguinal white adipose tissue (iWAT), #4 mammary fat pad, and brown adipose tissue (BAT) from WT (n=11-16) and L-KO (n=13-16) mice. (G) Annotated serum biomarkers of liver damage in WT (n=6) and L-KO (n=9) mice. (H) Representative H&E-stained histochemical images of the liver from male WT (top) and Gclc L-KO (bottom) mice (scale bars = 200 µm). (I-J) Gene Set Enrichment Analysis of (I) upregulated oxidative stress response genes and (J) downregulated lipogenic-related genes in the liver of L-KO mice compared to WT mice. (K) Relative expression of Hmox1 mRNA in the liver of WT (n=4-8) and L-KO (n=4-7) mice following treatment with AAV-TBG-Cre. (L) Relative expression of NQO1 protein in the liver of WT and L-KO mice. Expression of actin was used as a control. (M) Relative abundance of annotated lipid precursors in the liver of WT (n=4) and L-KO (n=4) mice three weeks following treatment with AAV-TBG-Cre. Data are shown as mean ±SEM. An unpaired two-tailed t-test was used for (A-G). A one-way ANOVA with subsequent Tukey’s multiple comparisons test was used for (K) and (M) to determine statistical significance. ns = not significant, * P value < 0.05, ** P value < 0.01, *** P value < 0.001, **** P value < 0.0001.

**Figure S5.**
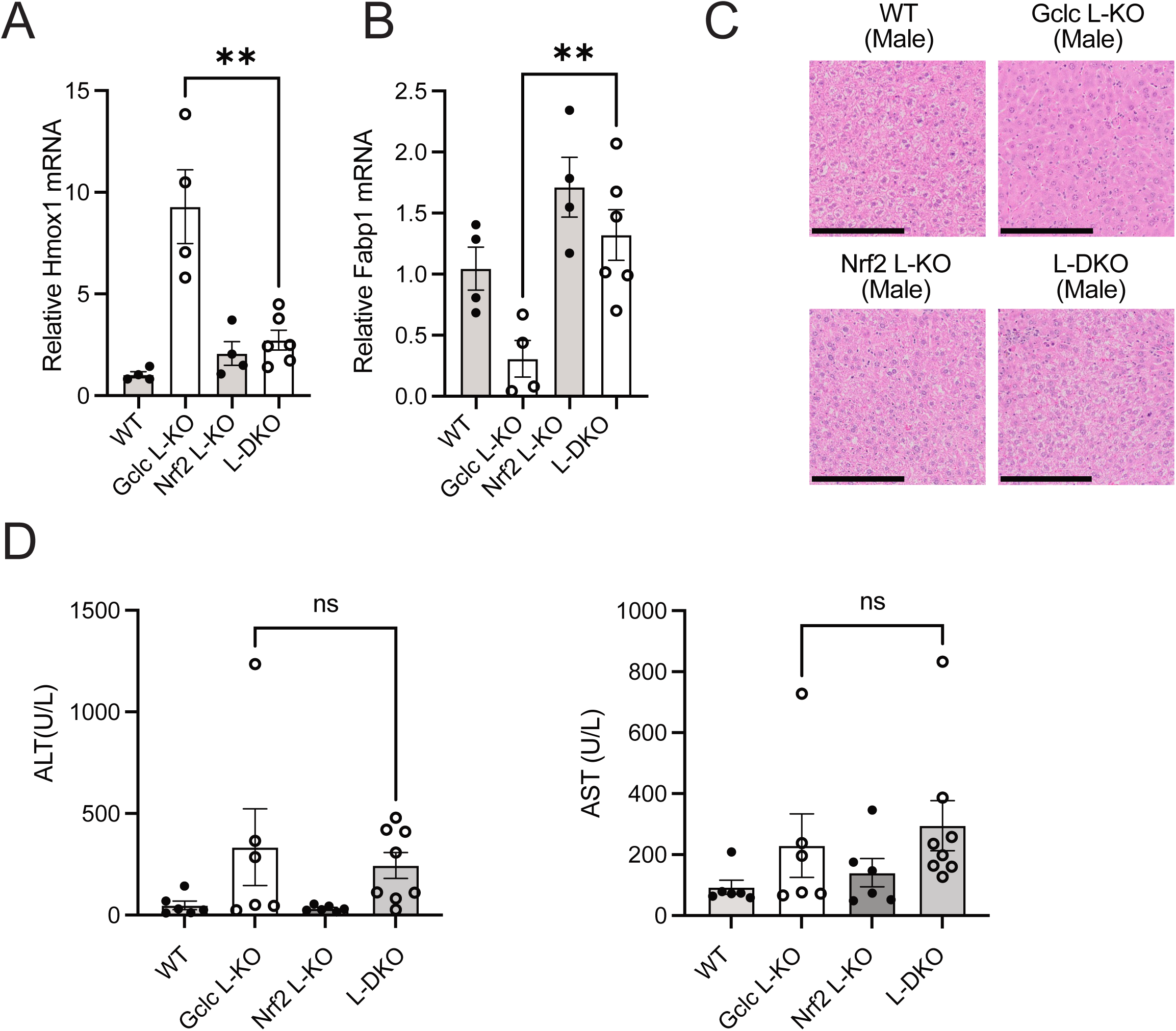
*Nrf2* deletion in liver Gclc- KO mice does not cause liver damage. Related to Figure 5. (A-B) Relative mRNA expression of (a) Hmox1 and (b) Fabp1 in the liver of WT (n=4), Gclc L-KO (n=4), Nrf2 L-KO (n=4), and L-DKO (n=6) mice. Expression levels were normalized to the expression of the reference gene Rps9. (C) Representative H&E-stained histochemical images of the liver from male Gclc WT, Gclc L-KO, Nrf2 L-KO, and L-DKO mice (scale bars = 200 µm). (D) Concentration of annotated serum biomarkers indicative of liver damage in WT (n=6), Gclc L- KO (n=6), Nrf2 L-KO (n=6), and L-DKO (n=8) mice. Data are shown as mean ±SEM. A one-way ANOVA with subsequent Tukey’s multiple comparisons test was used for (A-B). An unpaired two-tailed t-test was used for (D) to determine statistical significance. ns = not significant, * P value < 0.05, ** P value < 0.01, *** P value < 0.001, **** P value < 0.0001.

**Supplementary Figure 6.**
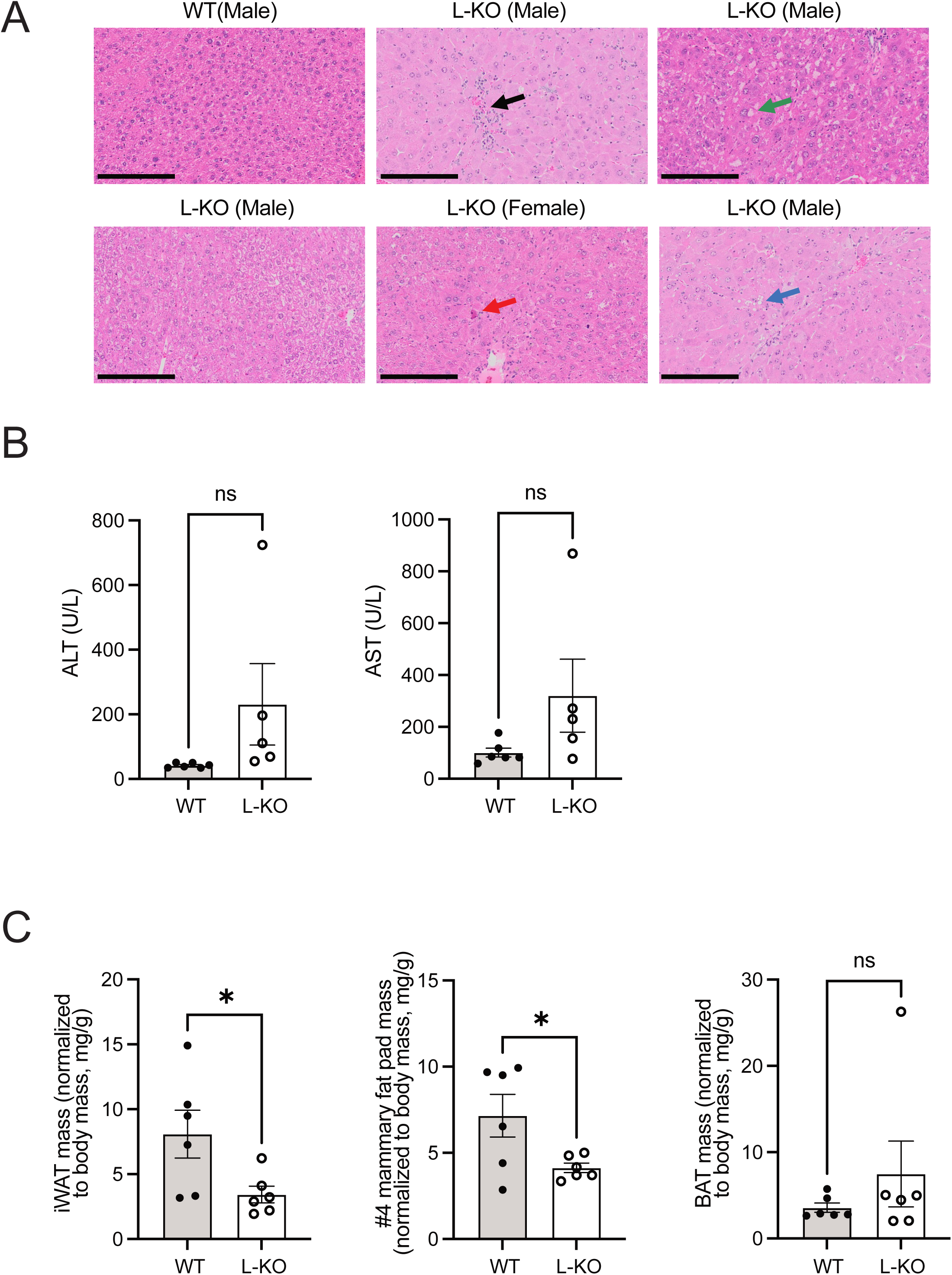
Prolonged liver-specific *Gclc* deletion induces fat tissue loss. Related to Figure 6. (A) Representative H&E-stained histochemical images of liver from WT (n=1) and Gclc L-KO (n=5) mice (scale bars = 200 µm). The black arrow indicates leucocyte infiltration; the green arrow indicates possible incipient balloon changes; the blue arrow indicates incipient steatosis; the red arrow indicates an apoptotic hepatocyte (B) Concentration of serum biomarkers of liver damage in WT (n=6) and L-KO (n=5) mice. (C) Mass of adipose tissue depots normalized to body mass; inguinal white adipose tissue (iWAT) (left), 4^th^ mammary fat pad (middle), and brown adipose tissue (BAT) in WT (n=6) and L-KO (n=6) mice. Data are shown as mean ±SEM. An unpaired two-tailed t-test was used for (B-C) to determine statistical significance. ns = not significant, * P value < 0.05, ** P value < 0.01, *** P value < 0.001, **** P value < 0.0001.

